# Agency and Expectations in Pain Treatment: An Investigation of the Active Inference Model

**DOI:** 10.1101/2022.07.20.500796

**Authors:** Andreas Strube, Björn Horing, Michael Rose, Christian Büchel

## Abstract

The fact that we cannot tickle ourselves is traditionally explained by the attenuation of somatosensation by predictions from a forward model of self-generated movements. Alternatively, it has been suggested within the framework of active inference that Bayes-optimal reduction of sensory precision can explain this phenomenon. Combining a pain paradigm with stimulus-related expectations allows to compare both models through predictions from the Bayesian account of expectation-based hypoalgesia, where pain is influenced by the precisions of somatosensation and expectation. In two experiments, heat pain was sham-treated either externally or by the subject, while a cue created higher or lower treatment expectations. Both experiments revealed greater pain relief under self-treatment and high treatment expectations. Electroencephalography revealed a modulation of theta-to-beta frequencies linked to agency and top-down modulations of pain perception. Computational modeling showed that this is better explained by an attenuation of somatosensation than a downregulation of somatosensation precision, favoring the forward model.

## Introduction

Charles Darwin observed that “from the fact that a child can hardly tickle itself, or in a much less degree than when tickled by another person, it seems that the precise point to be touched must not be known”^1^. The phenomenon that self-generated touch feels less intense – and less ticklish – than identical externally generated touch has been reproduced repeatedly^2–5^. In the classical formulation of the forward model^6^, this somatosensory attenuation is explained by continuously generated predictions of the sensory consequences of a motor command and these accurate predictions are used to attenuate the sensory consequences of self-produced movement. Under this model, smaller prediction errors during self-generated movement lead to a percept of a less intense sensation, relative to externally generated unpredicted outcomes.

An alternative explanation of sensory attenuation has been proposed in the framework of active inference^7^. In active inference^8,9^, the brain consistently attempts to reduce prediction errors of its generative model of the world. It is proposed that prediction errors can be minimized in one of two ways. As a first option, we can minimize prediction errors by changing predictions to explain sensory input (i.e. we update our model in a way that explains the world better). Alternatively, we can minimize prediction errors by performing an action (i.e. we act in a way that the world fulfills our predictions). In order to act, a contradiction has to be resolved: Self-generated actions (which are proprioceptive predictions to move) must overcome sensory information of the current state of the world – namely, the sensory information that we are currently not moving. Proprioceptive predictions to move would therefore be inhibited by prediction errors elicited by sensory information that one is currently not moving. This contradiction can be resolved by lowering the precision of sensory evidence to the consequences of one’s own actions^7^.

To disentangle predictions derived from active inference from the classical forward model, the paradigm of expectation-based hypoalgesia can be helpful. Placebo hypoalgesia is seen as an expectancy-driven phenomenon, which describes a pain relief mediated by prior experiences, in the absence of active treatment^10–14^. It has been proposed that pain perception can be seen as a Bayesian problem requiring the integration of expectations with stimulus intensity^15–18^. Expectations are integrated with incoming nociceptive stimulus information, and both are weighted by their respective precision to form a pain percept. This has been shown by manipulation of the level of precision of prior treatment expectations, where expectation-based effects were more pronounced with more precise treatment expectations^19^.

Similar to an application in visual perception^20^, less precise information about the current stimulation would lead to a relatively higher influence of prior expectation, while more precise stimulus information would lead to less influence of prior expectation on perception. Importantly, it is possible to design a placebo hypoalgesia paradigm in which sensory evidence (i.e. enhanced or reduced treatment efficacy) is either self-generated or externally generated. The active inference framework predicts that self-treatment would lead to a reduced precision of somatosensation and consequently to a lower weight of somatosensation; therefore, positive and negative expectations would have a relatively large impact on pain intensity in the respective direction. In contrast, the forward model predicts that self-treatment should always lead to a decrease in perceived intensity; therefore, self-treatment would always be associated with a lower pain percept, compared to external treatment, regardless of whether a positive or negative expectation existed.

It has been shown that pain perception is modulated by agency on a neurophysiological and behavioral level^21–35^. This beneficial effect is utilized in Patient-Controlled Analgesia (PCA) commonly used in post-operative care – patients receiving PCA experience less pain as compared to patients receiving traditional (i.e. externally applied) analgesia^36,37^.

In two experiments, we manipulated both expectations and agency. We established the illusion that heat pain could be treated through two variants of (putative) TENS (Transcutaneous Electric Nerve Stimulation) associated with either a highly effective or less effective treatment effect. In reality, treatment was implemented by reducing the temperature of the heat stimulator. This supposed treatment was either actively started by the participant (self-treatment) or by the experimenter (external treatment, actually administered by computer). In an initial stage, participants were conditioned to expect different treatment outcomes by an actually stronger reduction of the painful heat stimulation, as predicted by a visual cue indicating highly effective treatment. During the test stage following the conditioning, treatment was identical, irrespective of the cue indicating either highly effective or less effective treatment (i.e. heat was reduced to the same level in both conditions). In this fashion, we combined placebo (expectation is better than the test stimulus) and nocebo (expectation is worse than the test stimulus) expectation effects in pain treatment, creating bidirectional modulations of pain by placebo/nocebo expectations.

Derived from the Bayesian model of expectation-based hypoalgesia (see Fig. 1), we hypothesized that if the forward model applied, self-treatment would result in overall lower pain ratings, i.e. regardless of prior expectations. In contrast, if the active inference model applied, less precision of self-generated sensory consequences would enhance expectation effects (i.e. placebo/nocebo effects); this would manifest in pain ratings influenced more strongly by expectations in self-treatment test trials. See Fig. 1d for statistical hypotheses based on the forward model; see Fig. 1e for statistical hypotheses based on active inference.

**Figure 1.**
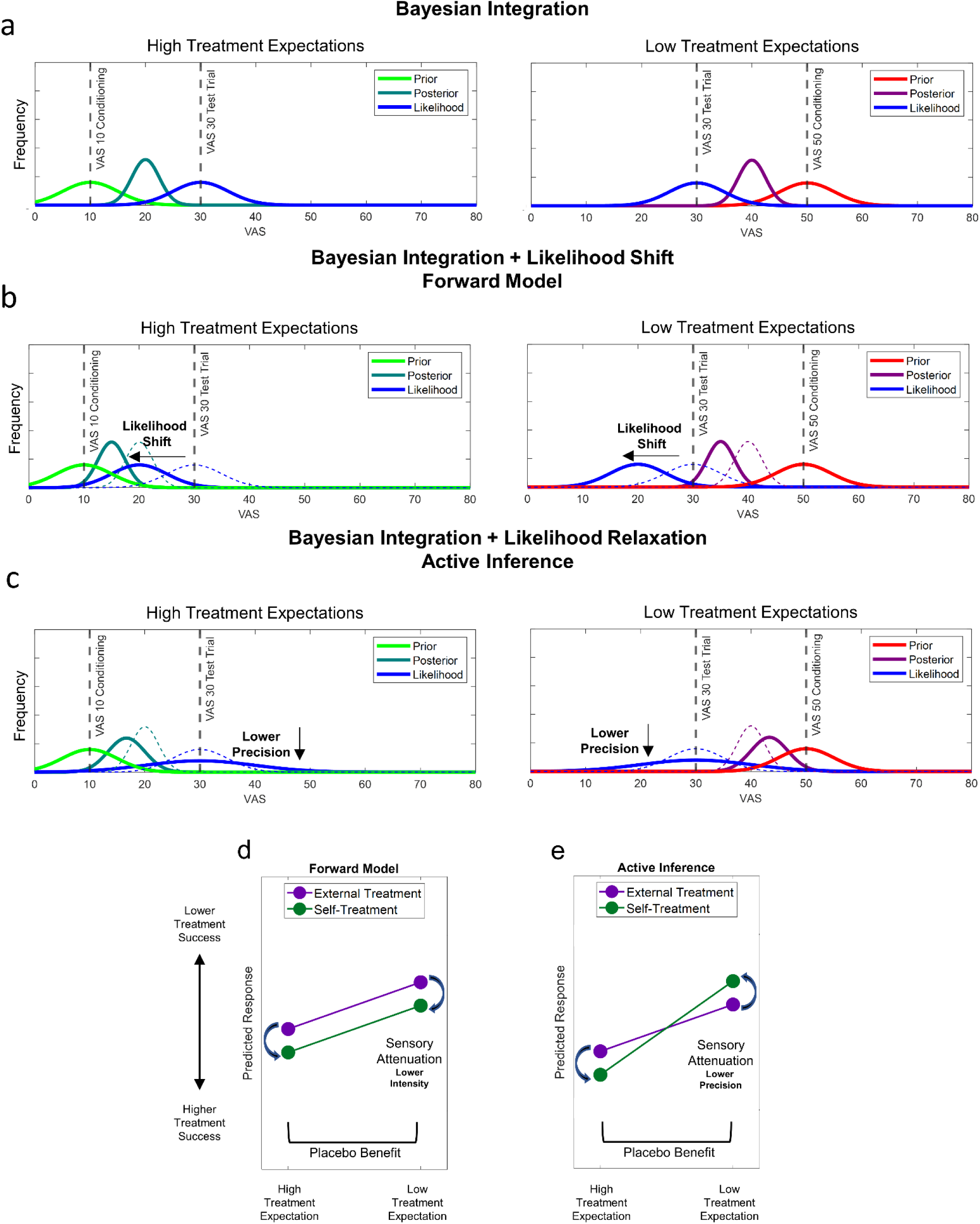
Bayesian models of pain perception. Bayesian model comparison was used to evaluate two Bayesian pain placebo/nocebo models. (a) The core of both models is the Bayes-optimal integration of prior experiences (here centered at VAS = 10 for placebo and at VAS = 50 for nocebo) with incoming nociceptive information (i.e. likelihood) to form a percept (i.e. posterior). Prior, likelihood and posterior were approximated by Gaussian distributions allowing for an analytic solution of Bayesian integration (for details see supplemental methods). The sensory attenuation model (b) has a free parameter that allows to shift the likelihood mean for self-treatment trials (see Methods, Eq.2). For example, the likelihood mean for self-treatment can be shifted (blue, solid line) as compared to the mean for external treatment (blue, dotted line). In high treatment expectation (placebo), this will lead to a shift of the posterior (dark green, solid line) to lower VAS values as compared to external (dark green, dotted line) because of the integration of the shifted lower likelihood with the prior. Similarly, in low treatment expectation (nocebo), this will lead to a shift of the posterior (purple, solid line) to lower VAS values as compared to external treatment (purple, dotted line). This is in contrast to the active inference model (c) which has a free parameter that can change likelihood precision (see Methods, Eq.3). If self-treatment is linked to a lower likelihood precision (blue, solid line) as compared to external treatment (blue, dotted line) this model should explain the data better than the forward model. As an example, this can lead to a posterior (dark green, solid line) which is more strongly drawn to the prior (VAS10 conditioning) in self-treatment than external treatment, due to the lower “impact” of the likelihood (dark green, dotted line). In low treatment expectation, the posterior (purple, solid line) would be drawn more strongly to the prior (VAS50 conditioning) in self-treatment than external treatment (purple, dotted line). Note that for actual modeling we utilized individual prior and likelihood parameters, whereas here, parameters are based on calibration target values for illustration purposes. The forward model (d) predicts a decrease of perceived stimulus intensity in self-treatment (green line) as compared to external treatment (purple line), meaning a higher treatment success in self-treatment trials as compared to external treatment trials. In active inference (e), self-treatment is associated with a decrease in precision, and thus a larger influence of expectations in self-treatment (green line) as compared to external treatment (purple line).

In placebo test conditions, the forward model would predict a better treatment outcome due to sensory attenuation leading to lower sensed stimulus intensity in self-treatment (as compared to external treatment). The active inference model would also predict a stronger influence by expectations, but due to sensory attenuation leading to lower stimulus precision. Thus, in placebo conditions, both models predict an improvement of treatment outcome by self-treatment as compared to external treatment. In nocebo conditions (i.e. a worse outcome is expected than the actual outcome) on the other hand, the forward model would also suggest an improved treatment outcome in self-treatment, due to lower sensed stimulus intensity. In contrast, crucially, active inference would predict a stronger influence of nocebo expectations due to a lower stimulus precision in self-treatment and thus, relatively weaker treatment success in self-treatment as compared to external treatment. This can also be translated to formal models of Bayesian integration (Fig. 1a) in pain perception incorporating the forward model (Fig. 1b) and active inference (Fig. 1c), respectively. With these models, we performed a formal Bayesian model selection (see Fig. 1 for an overview of the candidate models and model predictions).

The first experiment (*N* = 25) used continuous pain ratings to establish a precise readout of pain perception during painful heat stimulation and after treatment, while the second experiment (*N* = 54) additionally employed electroencephalography (EEG) to further evaluate neurophysiological correlates of the top-down modulation via expectations and agency. In both experiments, we applied heat pain to capsaicin-sensitized skin on the left radial forearm, after individual calibration to create comparable pain levels for each participant. To avoid a contamination of EEG data by movement artifacts through button presses by continuous pain ratings, we altered the paradigm for experiment 2 to include single outcome ratings instead of the continuous pain rating (see Fig. 2 for a graphical overview of the trial design).

**Figure 2.**
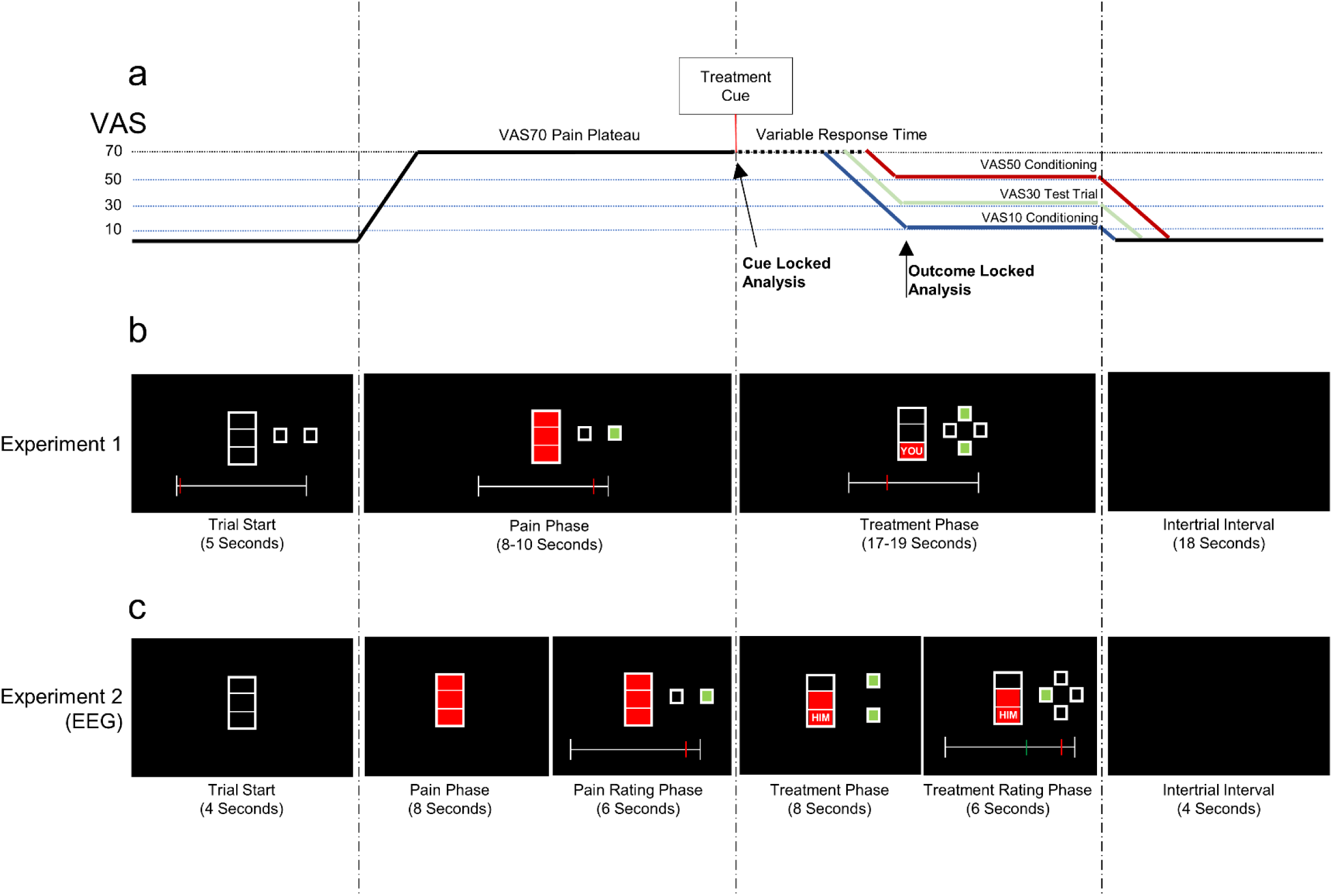
(a) Schematic representation of the paradigm, (b) trial design for experiment 1, and (c) trial design for experiment 2. Colored lines represent VAS50 conditioning (red), test trials (VAS30; green) and VAS10 conditioning (blue). The black line represents alterations in temperature common to all trial types, blue, green and red lines represent changes based on VAS10 conditioning, VAS30 test trials and VAS50 conditioning trials, respectively. At trial start, the thermal-heat stimulator (thermode), attached to the left radial forearm of the participant, is at the baseline temperature (set to 30°C for experiment 1 and to 28°C for experiment 2). A red bar indicates the start of the pain phase concurrent with an increase of thermode temperature to the individually calibrated pain level of VAS70. The start of the treatment phase is indicated by a cue showing whether self- or external treatment and whether highly or low effective treatment follows. This then leads to actual low (VAS10) or high (VAS50) temperatures during conditioning trials, respectively. In test trials, the final temperature is always at VAS30 regardless of the cued treatment efficacy. Arrows indicate time points for EEG data locks, i.e. the time axis of EEG time-frequency data was set to 0 according to the onset of the cue and to the treatment outcome (i.e. target treatment VAS level was reached by the thermode), respectively. In experiment 1 (b), a rating scale controlled with two buttons was presented during the whole trial. At trial start (5s), an empty bar was presented alongside the rating scale (set to 0 at the beginning) and a display of rating buttons (lighting up in green when pressed). During the following pain phase (8-10s), the empty bar was filled red as an indication for pain. After the pain phase, the treatment cue was presented. The treatment cue showed a reduction of the red bar, where a reduction to 1/3 of the total height was associated with highly effective treatment and a reduction to 2/3 of the total height was associated with a low effective treatment. Additionally, a signal word indicated self- or external treatment. After a lag of 2s, the treatment buttons appeared on the display, lighting up in green when pressed by the subject or externally. After the treatment button was pressed, the temperature was decreased to the respective pain level. An ITI (intertrial interval) of 18s followed. In experiment 2 (c), pain ratings scales and buttons were only presented during designated rating phases. Treatment could be started immediately after the onset of the treatment cue. Here, the timing of each trial was: trial start (4s), pain phase (8s), pain rating phase (6s), treatment phase (8s) and treatment rating phase (6s) with an ITI of 4s.

## Results

### Experiment 1: Behavioral results

The first experiment used continuous pain ratings (Fig. 3).

**Figure 3.**
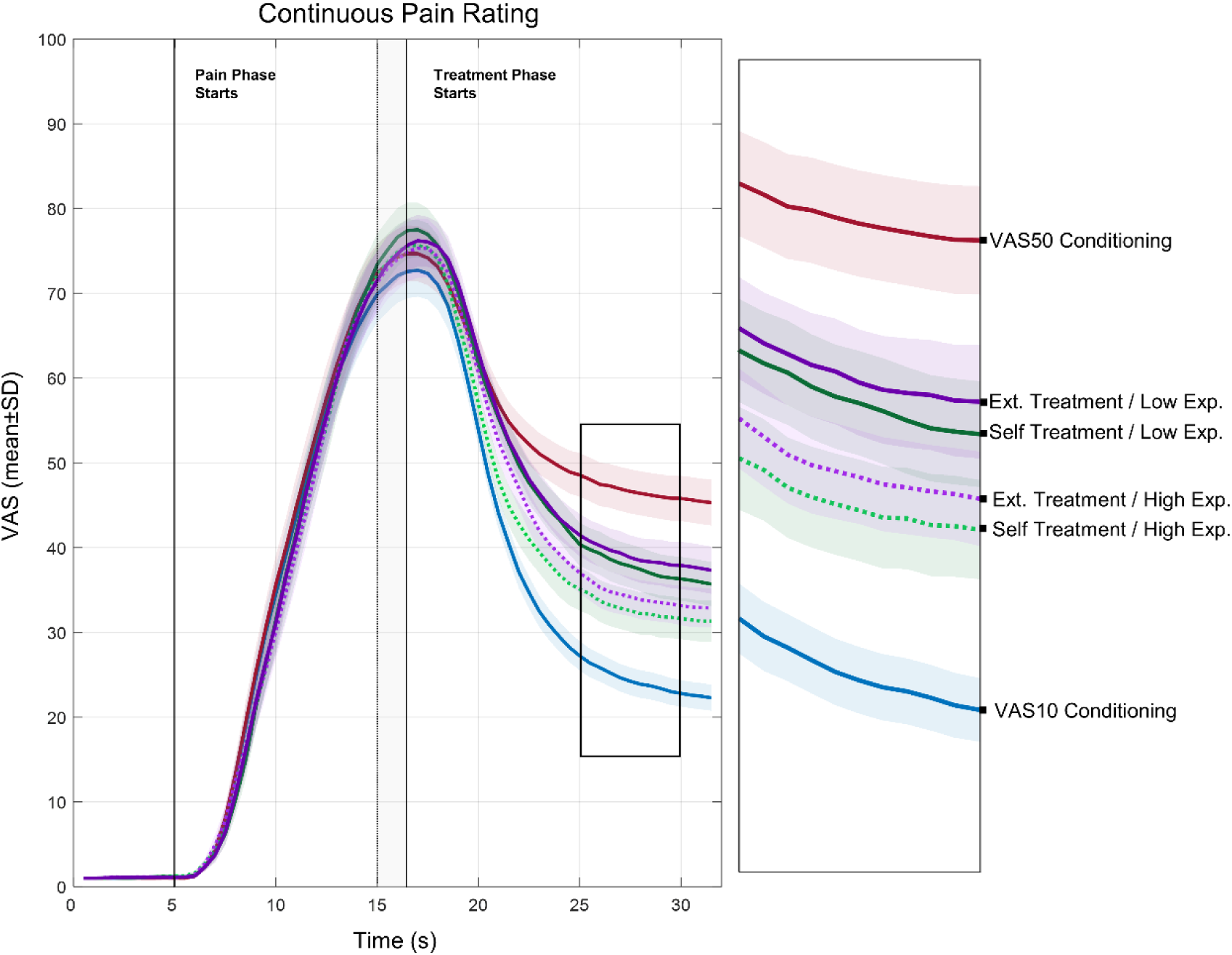
Continuous VAS ratings per condition. Each line represents a different condition, i.e. VAS10 and VAS50 conditioning, and 4 test conditions following VAS30 (self- versus external treatment, low versus high treatment expectation). Pain phase (VAS70) starts after a cue presentation of 5s for a jittered duration of 8-10s. Afterwards the treatment phase started, beginning with the presentation of the treatment cue for 2s. Then, treatment was started either by the participant or externally. Lines on the right represent an enlargement of the highlighted section (25-30s).

The treatment outcome differed objectively as we reduced the pain stimulus to three different intensities, i.e. VAS10 and VAS50 for high and low treatment efficacy respectively during conditioning trials, and VAS30 for test trials (albeit presented with the respective high or low conditioned cues). To evaluate if participants experienced these three intensities to be different, we conducted a repeated measures ANOVA on the final continuous rating data points (post-treatment VAS rating) from all three stimulus intensities, including conditioning and test trials (averaging across expectations) which revealed a significant difference (*F*(2,48) = 43.78, *p* < 0.001, *η_p_^2^* = 0.646) (Fig. 4a).

**Figure 4.**
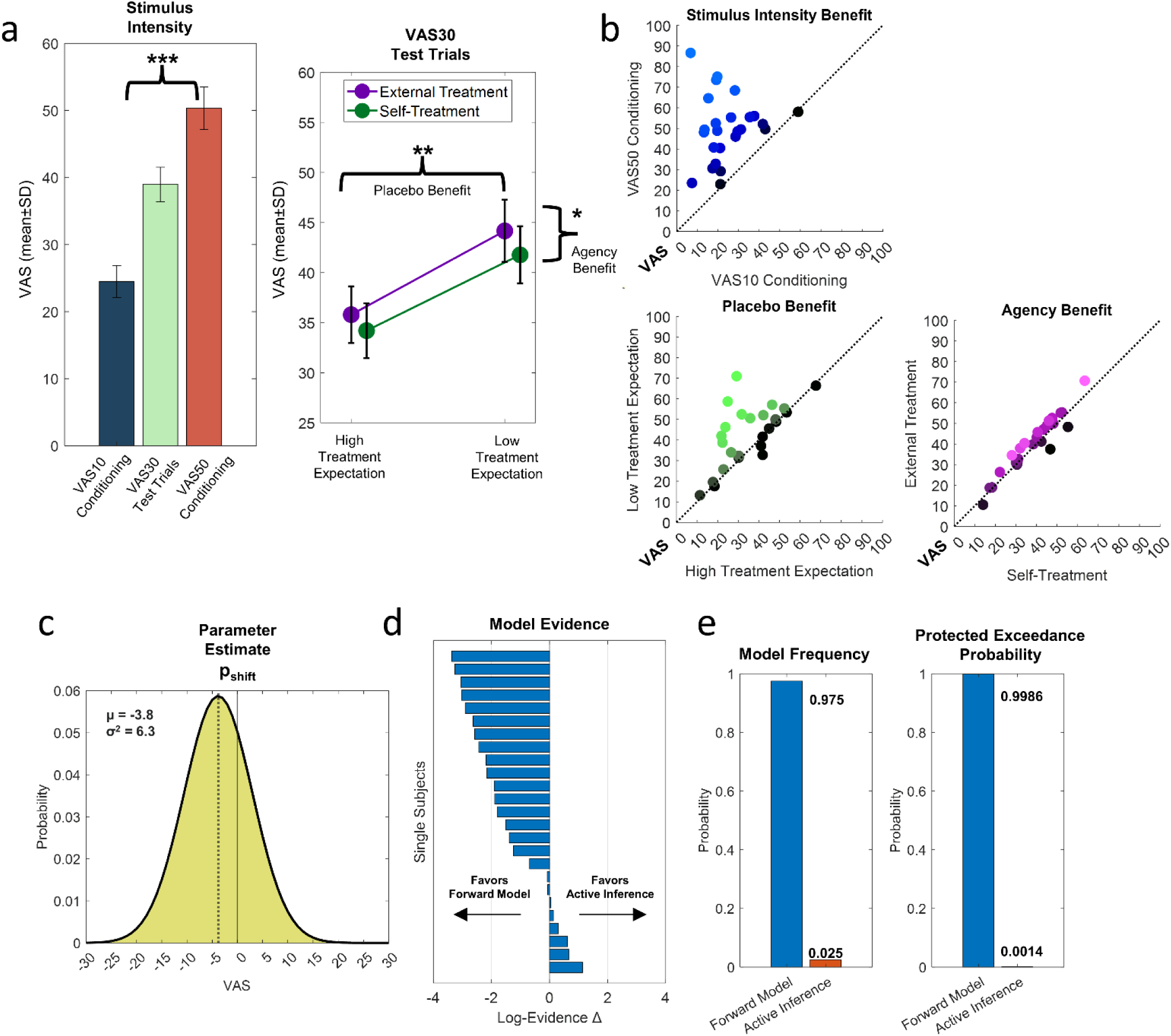
Results from VAS (Visual Analogue Scale) rating analyses of experiment 1 (*N* = 25). (a) Post-treatment VAS ratings for each stimulus intensity condition (VAS10, 30 and 50) and lines representing the contrasts of low versus high treatment expectation and self- versus external treatment during test trials. Bars and lines represent post-treatment VAS ratings averaged per condition. (b) Scatter plots represent single subject values for treatment outcomes for conditioning, expectation, and agency. Scatter plots represent contrasts of conditions, i.e. each dot represents averaged ratings of a single subject for VAS50 versus VAS10 conditioning (blue), low treatment expectation versus high treatment expectation (green) and external treatment versus self-treatment (purple). Brighter colors indicate larger benefits of stimulus intensity (VAS10 versus VAS50 conditioning), placebo benefits (high treatment expectations versus low treatment expectations) and agency benefits (self-treatment versus external treatment). Data points above the diagonal represent single subjects with stimulus intensity, placebo and agency benefits, respectively. (c) Probability density function of group parameter estimates for the likelihood shift parameter p_shift_ of the winning forward model. (d) Single subject differences of log evidence for the forward model versus active inference model (negative values favor the forward model) and (e) model frequencies and protected exceedance probabilities.

Post-hoc analyses using Bonferroni correction for multiple comparisons indicated that all three stimulus intensity levels differed significantly from each other, revealing higher post-treatment VAS ratings for VAS50 conditioning trials (*M* = 50.35, *SD* = 15.78) versus VAS30 test trials (*M* = 38.98, *SD* = 12.86) and for VAS30 test trials versus VAS10 conditioning trials (*M* = 24.47, *SD* = 11.95; all *p* < 0.001).

As a next step, we evaluated the effects of our manipulations for the test trials, where intensity of the painful stimulus was always reduced to an individually calibrated level of VAS30. Here, post-treatment VAS ratings could either be influenced by agency (self-versus external treatment), expectations (low versus high treatment expectations), or their interaction. Considering the forward model (see Fig. 1d), we would expect higher treatment success in self-treatment test trials as compared to external treatment test trials regardless of prior treatment expectations. In other words, we would expect a main effect of agency with or without a main effect of expectation, but no interaction between both factors. Conversely, for the active inference model (see Fig. 1e), by lowering the precision of sensory evidence, prior treatment expectations would gain a higher relative weight compared to the sensory information in self-treatment trials as compared to external treatment trials, which would manifest itself as an interaction (i.e. expectation effects should be larger in self-treatment).

Here, again we conducted a repeated measures ANOVA to test for main effects of agency, expectation, and their interaction in the test conditions (VAS30). We found a significant sensory attenuation effect, that is, a main effect of agency (*F*(1,24) = 6.2, *p* = 0.02, *η_p_^2^* = 0.205), meaning that post-treatment VAS ratings were lower for self-treatment trials (*M* = 37.99, *SD* = 12.57) versus external treatment trials (*M* = 39.98, *SD* = 13.45). Furthermore, we found a significant expectation effect (*F*(1,24) = 10.738, *p* = 0.003, *η_p_^2^* = 0.309), i.e. high treatment expectations were associated with lower post-treatment VAS ratings (*M* = 35.00, *SD* = 13.66; conditioned with VAS10) than those following low treatment expectations (*M* = 42.97, *SD* = 14.77; conditioned with VAS50). Importantly, we did not observe a significant interaction of expectation and agency (*F*(1,24) = 0.679, *p* = 0.42, *η_p_^2^* = 0.028).

For model-based analysis of our behavioral data, we created two Bayesian models of pain perception in placebo pain treatment (see Fig. 1) which were inverted and compared using variational Bayesian methods (VBA, see Methods for details). We used the Bayesian integration model of pain perception for both the active inference model and the forward model (Fig. 1a). In self-treatment test trials under the forward model, we included the parameter p_shift_ which allowed for a shift of the likelihood and thus for a posterior distribution which was shifted into the same direction in low and high treatment expectation conditions, as predicted by the forward model (Fig. 1b). This was contrasted to the active inference model, where the posterior distribution should be drawn to the conditioned pain experience which is explained by a decrease of the likelihood precision, and thus we included the parameter p_relax_ which allowed for such a decrease in likelihood precision (Fig. 1c).

We used a random effects (RFX) Bayesian model selection approach^38,39^ to estimate the overall posterior model probability across subjects. The RFX model exceedance probability was at *φ* = 0.9986 for the forward model compared to *φ* = 0.0014 for the active inference model. Hence, we see clear evidence for the forward model with a free parameter enabling likelihood shift over the active inference model with a free parameter allowing a relaxation of likelihood variance (see Fig. 4e). Both models (*φ* > 0.999) outperformed a null model without these free parameters (*φ* < 0.001), and a more complex full model including both free parameters (*φ* < 0.001). See Supplementary Fig. 1 for a comparison of all candidate models. See Supplementary Fig. 2 for forward model comparison with the null model and full model and Supplementary Fig. 3 for respective active inference comparisons.

In summary, results using continuous pain ratings clearly demonstrate both sensory attenuation effects (i.e. self-treatment lead to better outcomes) and expectation effects (i.e. high treatment expectations lead to better treatment outcomes), but no interaction between expectation and agency. Model selection provides strong evidence in favor of the forward model over the active inference model (see Fig. 4 for a summary of the results).

### Experiment 2: Behavioral results

As in the first experiment, a repeated measures ANOVA with all three stimulus intensities (including conditioning and test trials) revealed a significant difference (*F*(2,106) = 118.32, *p* < 0.001, *η_p_^2^* = 0.691) between all three intensities (VAS10, 30 and 50) in post-treatment VAS ratings. Post-hoc analyses using Bonferroni correction for multiple comparisons indicated that all three stimulus intensity levels differed significantly from each other, revealing higher post-treatment VAS ratings for VAS50 conditioning trials (*M* = 49.66, *SD* = 15.46) versus VAS30 test trials (*M* = 38.24, *SD* = 17.18) and for VAS30 test trials versus VAS10 conditioning trials (*M* = 31.04, *SD* = 17.45; all *p* < 0.001).

For the evaluation of main effects of agency and expectation and their interaction for the test trials, we again conducted a repeated measures ANOVA. We observed a significant sensory attenuation effect (*F*(1,53) = 19.13, *p* < 0.001, *η_p_^2^* = 0.265), i.e. post-treatment VAS ratings were lower for self-treatment trials (*M* = 37.15, *SD* = 17.26) as compared to external treatment trials (*M* = 39.33, *SD* = 17.30). Also, we found a significant expectation effect (*F*(1,53) = 35.57, *p* < 0.001, *η_p_^2^* = 0.402), i.e. high treatment expectations (*M* = 34.91, *SD* = 17.69; conditioned with VAS10) were associated with lower post-treatment VAS ratings than low treatment expectations (*M* = 41.56, *SD* = 17.64; conditioned with VAS50). As in the first experiment, we did not observe a significant interaction of treatment expectation and agency (*F*(1,54) = 0.02, *p* = 0.887, *η_p_^2^* = 0.003).

Again, we used a random effects (RFX) Bayesian model selection approach to estimate the overall posterior model probability across subjects for the post-treatment VAS ratings in experiment 2. For experiment 2, the RFX exceedance probability of *φ* = 0.9995 for the forward model compared to *φ* = 0.0005 for the active inference model again strongly favored the forward model over the active inference model (Fig. 5e). Both models (*φ* > 0.999) outperformed a null model without these free parameters (*φ* < 0.001), and a full model including both free parameters (*φ* < 0.001). See Supplementary Fig. 4 for a comparison of all candidate models. See Supplementary Fig. 5 for forward model comparison with the null model and full model and Supplementary Fig. 6 for respective active inference comparisons.

**Figure 5.**
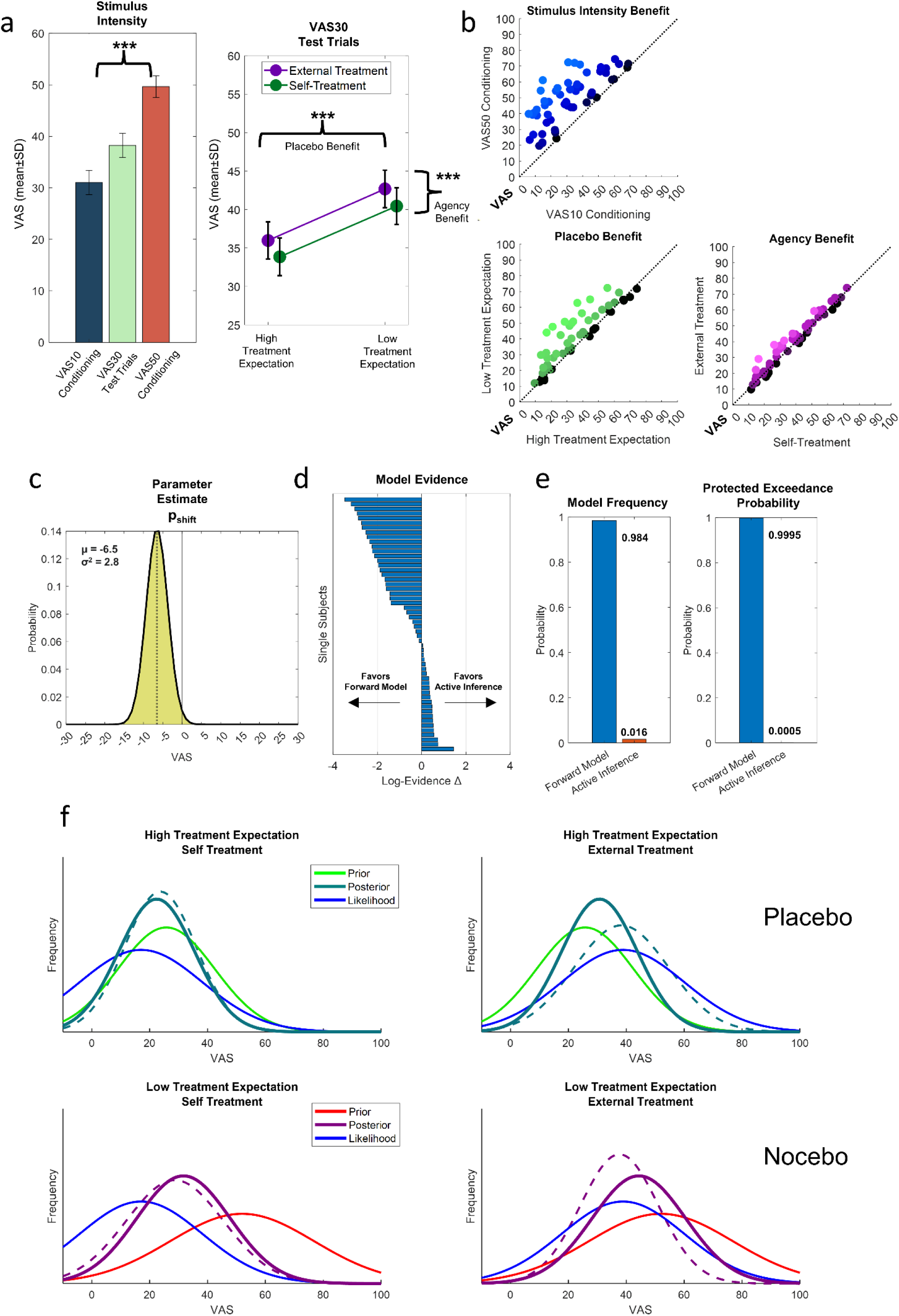
Behavioral VAS (Visual Analog Scale) post-treatment pain rating data of experiment 2 (*N* = 54). (a) Post-treatment VAS ratings for each stimulus intensity condition (VAS10, 30 and 50) and lines representing the contrasts of low versus high treatment expectation and self- versus external treatment during test trials. Bars and lines represent post-treatment VAS ratings averaged per condition. (b) Scatter plots represent single subject values for treatment outcomes for conditioning, expectation, and agency. Scatter plots represent contrasts of conditions, i.e. each dot represents averaged post-treatment VAS ratings of a single subject for VAS50 versus VAS10 conditioning (blue), low treatment expectation versus high treatment expectation (green) and external treatment versus self-treatment (purple). Brighter colors indicate larger benefits of stimulus intensity (VAS10 versus VAS50 conditioning), placebo benefits (high treatment expectations versus low treatment expectations) and agency benefits (self-treatment versus external treatment). Data points above the diagonal represent single subjects with stimulus intensity, placebo and agency benefits, respectively. (c) Probability density function of group parameter estimates for the likelihood shift parameter p_shift_ of the winning forward model. (d) Single subject differences of log evidence for the forward model versus active inference model (negative values favor the forward model) and (e) model frequencies and protected exceedance probabilities. (f) Example data of a single subject is shown for all test conditions, i.e. high treatment expectations with self- and external treatment and low treatment expectations with self- and external treatment. Lines represent empirical Gaussian high treatment expectation priors (green), low treatment expectation priors (red) and likelihood (blue) based on the fitted forward model. The solid dark green line represents the theoretical posterior based on Bayesian integration of the high treatment expectation prior and the likelihood. The dotted dark green line represents the parameters of a fitted Gaussian distribution to the empirical post-treatment VAS ratings of the respective conditions. Accordingly, the solid purple line represents the theoretical posterior based on Bayesian integration of the low treatment expectation prior and the likelihood. The dotted purple line represents the parameters of a fitted Gaussian distribution to the empirical post-treatment VAS ratings of the respective conditions.

Taken together, in the second experiment, we replicated the rating-related results of the first experiment. Again, these results demonstrate sensory attenuation effects (i.e. self-treatment leads to better treatment outcomes) and expectation effects (i.e. high treatment expectations lead to better treatment outcomes) but no interaction between expectation and agency. Model selection provides strong evidence in favor of the forward model over the active inference model (see Fig. 5 for a summary of the results).

### Experiment 2: EEG time-frequency data

For the statistical analysis of EEG data, we considered two separate time-points for time-frequency data to evaluate cue-locked as well as treatment outcome-locked effects. For cue-locked data, we set t = 0 to the onset of the cue indicating the conditioned effectiveness of the treatment and the agency condition of the treatment phase. Outcome-locked data sets t = 0 to the point when the thermode reached the calibrated treatment VAS target, and thus, takes individual variations in treatment latency into account. All tests were corrected for multiple comparison using Monte Carlo cluster tests (see Methods for details). See Supplementary Fig. 7-9 for z-scored cue-locked time-frequency data and Supplementary Fig. 10-12 for z-scored outcome-locked time-frequency data for all conditions at Fz, Cz and Pz, respectively.

### Conditioning

In each trial, a cue indicated an upcoming highly or low effective treatment on the ongoing VAS70 stimulus. This association was established during conditioning trials, where a cue indicating high treatment efficacy was associated with a (physical) stimulus intensity decrease to a temperature individually representing VAS10. Similarly, a cue indicating low treatment efficacy was associated with a decrease to a temperature representing VAS50. A cluster-corrected dependent samples t-test on cue-locked data (0 to 2s after the cue, 4-181Hz) revealed no differences between VAS10 and VAS50 conditioning trials immediately after cue presentation (all *p* > .05). However, differences were significant for outcome-locked data (in a window of -1 to 2s with t = 0 at target temperature, 4-181Hz), revealing three clusters of activity associated with different conditioning types (VAS10 versus VAS50; Fig. 6; also see Supplementary Fig. 13 for topographies of significant clusters of activity for different frequency bands and Supplementary Fig. 14 for bar graphs representing the averaged power at significant clusters of activity for all conditions). Negative times reflect activity during unfolding pain relief, whereas positive times indicate activity during the outcome phase, i.e. when temperatures were stable at the final outcome level. We found a positive cluster (*p* < .001) including frequencies from 7 to 75Hz in a time frame from -0.4 to 1s, indicating an increase of EEG power for VAS10 conditioning versus VAS50 conditioning. Additionally, we found two negative clusters associated with lower EEG power of VAS10 versus VAS50 conditioning. Firstly, we found a negative cluster (*p* = 0.01) in the theta range including frequencies from 4 to 9.5Hz in a time frame from -0.65 to 0.9s. Secondly, we found another negative cluster (*p* = 0.043) ranging from -1 to -0.35s including frequencies from 9.5 to 45Hz. This demonstrates a modulation of theta (4-7Hz), alpha-to-beta (8-30Hz) and low gamma (31-50Hz) frequencies by treatment efficacy via differences in stimulus intensity, i.e. more effective treatment (VAS10) was associated with lower alpha-to-beta activity during the relief phase followed by lower theta and increased alpha-to-beta power at the treatment outcome phase, as compared to less effective treatment (VAS50).

**Figure 6.**
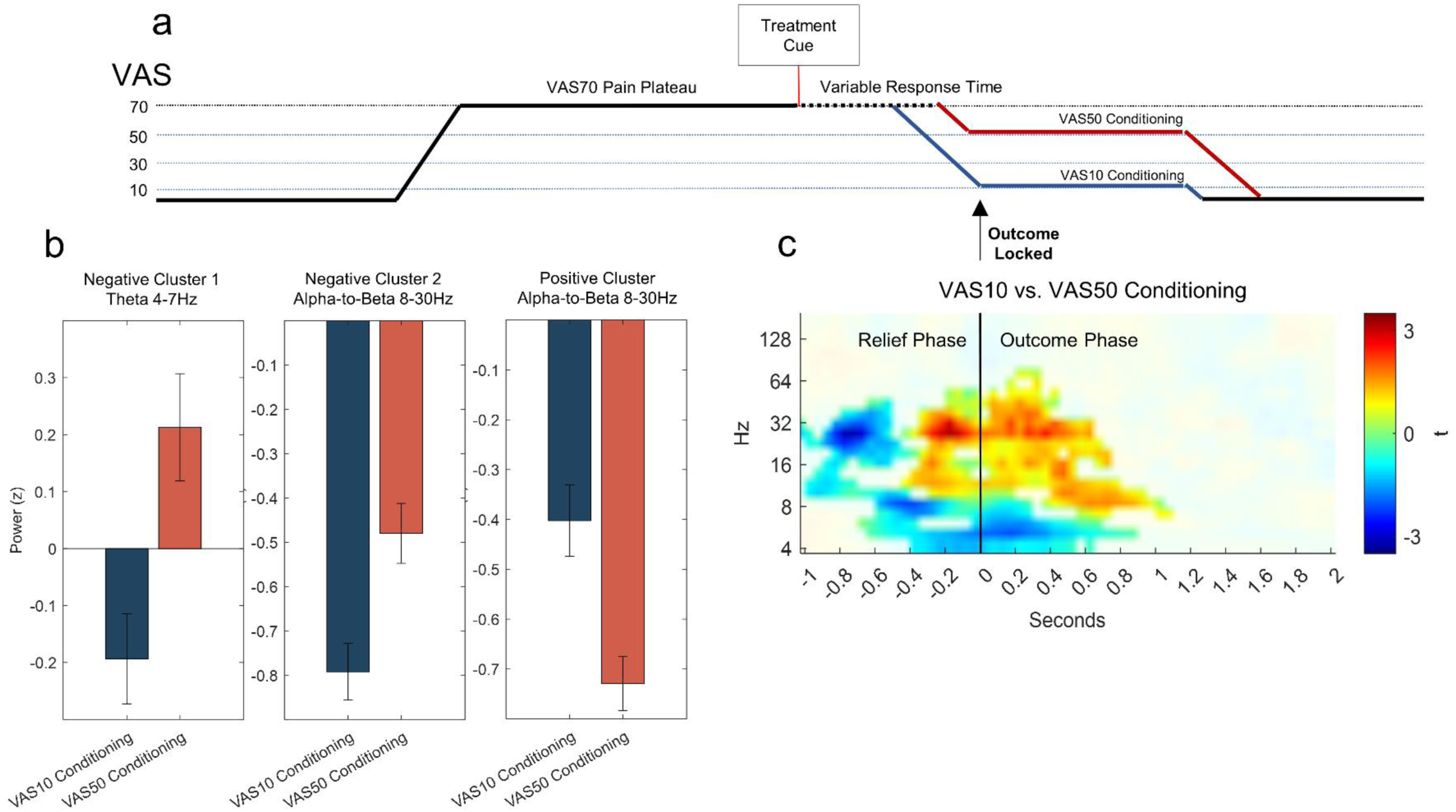
(a) Pain treatment paradigm, (b) differences in EEG power (z-scored) for VAS10 conditioning and VAS50 conditioning averaged over all significant samples in the theta (4-7Hz) and alpha-to-beta (8-30Hz) bands, (c) time-frequency plot of significant clusters of activity associated with the outcome-locked main effects of VAS10 versus VAS50 conditioning. Time-frequency plots are averaged t-values over all channels including significant data points of the respective clusters of activity. Non-significant data points are masked out. Colors represent *t*-values.

### Agency

Agency was manipulated as either participants initiated the treatment themselves, or the treatment was initiated (putatively) by the experimenter. To test for the effects of agency, we again considered both phases and conducted t-tests contrasting self-treatment and external treatment during test trials. Here, a cluster-corrected dependent samples t-test revealed no association between EEG time-frequency data and agency with cue-locked data (0 to 2s after cue onset, 4-181Hz; all *p* > .05). Treatment outcome-locked data (−1 to 2s at target temperature, 4-181Hz) revealed two significant clusters of activity associated with agency (Fig. 7). Data showed a negative cluster (*p* < 0.001) ranging from -1 to 1s including frequencies from 4 to 45Hz, indicating a negative association between agency and low frequency EEG power during the relief phase (−1 to 0s) and the outcome phase (0 to 2s). Another positive cluster was significant (*p* = .028), ranging from -0.2 to 0.65s including frequencies from 9.5 to 54Hz, indicating higher alpha-to-beta (8-30Hz) and low gamma (31-50Hz) power during the treatment outcome phase for self-treatment versus external treatment.

**Figure 7.**
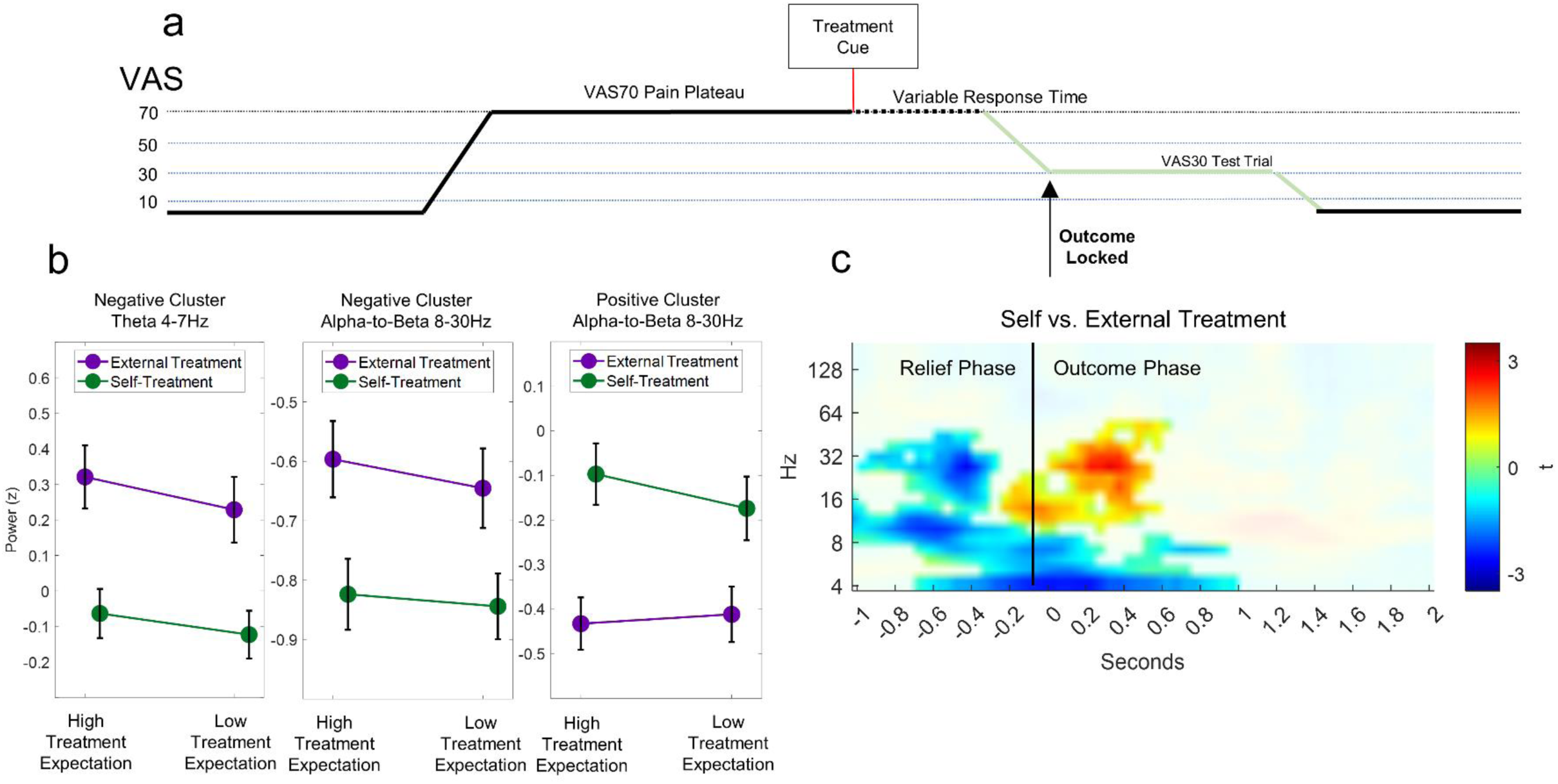
(a) Pain treatment paradigm, (b) a panel of line graphs representing differences in EEG power (z-scored) for self-treatment (green) versus external treatment (purple) averaged over all significant samples in the theta (4-7Hz) and alpha-to-beta (8-30Hz) bands, and (c) a time-frequency plot representing significant clusters of activity associated with the outcome-locked main effects of self- versus external treatment. Time-frequency plots are averaged t-values over all channels including significant data points of the respective clusters of activity. Non-significant data points are masked out. Colors represent *t*-values.

To further evaluate the influence of between-subject differences in sensory attenuation, we conducted a Pearson’s correlation analysis based on the Bayesian parameter estimate of the individually estimated likelihood shift parameter per subject (i.e. self-treatment benefit; see Methods for details). Here, no significant clusters were revealed associating between-subject differences in likelihood shift (i.e. self-treatment benefits) with EEG time-frequency data (all *p* > .05) in cue-locked (0 to 2s after cue onset, 4-181Hz) and treatment outcome-locked (−1 to 2s at target temperature, 4-181Hz) data.

The analysis of agency revealed two clusters associated with differences between self- and external treatment. Firstly, we observed a negative cluster during the relief and treatment outcome phase including frequencies from the theta (4-7Hz) to low gamma (31-50Hz) range. Secondly, when the target temperature was reached, data showed an increase of alpha-to-beta and low gamma activity associated with self-treatment (as compared to external treatment).

### Expectation

As a next step, we evaluated the effects of treatment expectations. During test trials, stimulus intensity (i.e. the target temperature of the treatment) was set to an individual value of VAS30. To evaluate the effects of different treatment expectations, we conducted a cluster-corrected dependent samples t-test on cue-locked data (0 to 2s after cue onset, 4-181Hz) which revealed no differences between high and low treatment expectations (all *p* > .05). Likewise, a cluster-corrected dependent samples t-test did not reveal significant clusters associated with treatment expectations at the treatment outcome (−1 to 2s at target temperature, 4-181Hz; all *p* > .05).

To further evaluate the effects of treatment expectations, we conducted a Pearson’s correlation on z-standardized behavioral expectation effects per subject (see Methods for details). Here, positive or negative clusters would indicate a correlative association of EEG power and the size of the expectation effects. However, no significant clusters were revealed associating between-subject expectation effects with EEG time-frequency data (all *p* > .05) in cue-locked (0 to 2s after cue onset, 4-181Hz) and treatment outcome-locked (−1 to 2s at target temperature, 4-181Hz) data.

### Interaction of agency and expectation

Finally, we tested for an interaction of treatment expectation and agency. At cue-locked data (0 to 2s after cue onset, 4-181Hz), a cluster-corrected dependent samples t-tests revealed a significant association of EEG power and the interaction term (*p* = .038; Fig. 8) ranging from 0 to 1.6s after cue onset and including frequencies from 4 to 13.5Hz. Here, we also conducted post-hoc t-tests, which revealed a crossed interaction (all *p* < .05, see Supplementary Information for detailed post-hoc t-test results confirming the crossed interaction). Treatment outcome-locked data did not reveal any cluster associated with an interaction of treatment expectation and agency (all *p* > .05).

**Figure 8.**
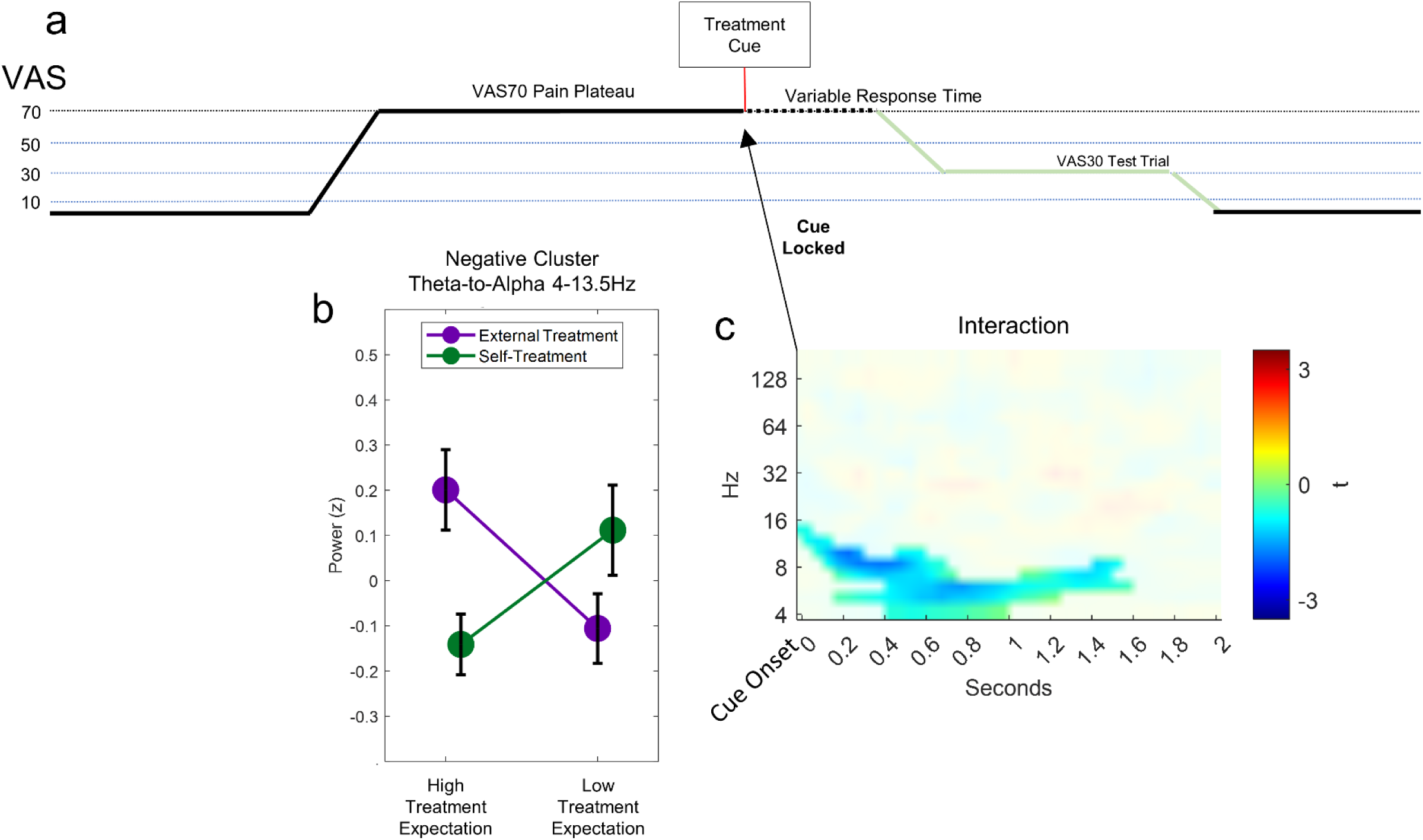
(a) Schematic graphical representation of the pain treatment paradigm, a line plot showing EEG power averaged over the significant agency x expectation interaction cluster for self-treatment (green) and external treatment (purple), (b) showing a significant crossed interaction, and (c) a time-frequency plot representing significant clusters of activity associated with the cue-locked interaction (agency x expectation) cluster. Time-frequency plots are averaged t-values over all channels including significant data points of the respective clusters of activity. Not significant data points are masked out. Colors represent *t*-values.

The analysis of the interaction term revealed differential integration of treatment expectation and agency information in theta (4-7Hz) and alpha (8-12Hz) frequencies starting shortly after the presentation of the cue (see Fig. 8 for a summary of the results).

## Discussion

In the present study, we demonstrated greater pain relief of self-treatment and high treatment expectations but no modulation of self-treatment benefits by expectations. Bayesian model selection provided strong evidence for the forward model (decrease of perceived intensity of sensory consequences of self-generated outcomes) over the active inference model (decrease of precision of sensory consequences of self-generated outcomes). These effects also manifested in EEG data: Stimulus intensity and agency modulated low frequency oscillatory responses in the theta (4-7Hz), alpha (8-12Hz), beta (13-30Hz) and low gamma (31-50Hz) ranges at the treatment outcome. Cue-related theta-to-alpha (4-12Hz) activity was differentially modulated by agency and expectation (representing a crossed interaction of agency and expectation).

We took great care to match both conditions (self- versus external treatment) with respect to cognitive and motor demands and adjusted the trials for visual, cognitive, and motor components. During both self-treatment and external treatment, a single button press was made by the subject - either to start the treatment or to acknowledge the (supposed) experimenter’s treatment. In addition, the correspondingly pressed buttons lit up on the screen, i.e. in both conditions two buttons changed to green, as in external treatment, self-treatment was “acknowledged” by the experimenter.

Concerning the role of agency, we expected an increased treatment efficacy^27,35^. Indeed, data from both experiments support this hypothesis: self-treatment was associated with lower post-treatment VAS ratings as compared to external treatment, despite identical objective stimulus intensity. Based on previous studies, we also expected that pain perception is modulated by expectation^40–45^. Both experiments support this hypothesis and showed a graded effect of expectation, i.e. high treatment expectations were associated with a higher treatment success than low treatment expectations.

Crucially, our experiment was designed to investigate the mechanism underlying improved treatment efficacy when treatment was self-initiated. The forward model^6^ posits that small prediction errors during self-generated movement lead to a percept of a less intense sensation, relative to externally generated unpredicted outcomes; applied to our pain protocol, this suggests an improvement in treatment outcome by self-treatment (see Fig. 1d). In contrast, the active inference model suggests that a decrease in precision of sensory consequences underlies the sensory attenuation phenomenon^7^. In a Bayesian sense, pain perception can be seen as the integration of expectation (prior) and nociceptive input (likelihood), with the precision of each term determining the amount of its contribution. Thus, as a consequence of the reduced nociceptive (likelihood) precision in the active inference model during our experiment, the effect of expectation should increase, as its precision remains constant. Consequently, we expected that the relative influence of prior expectations would be increased relative to the sensory evidence, which would be attenuated in precision. Therefore, self-treatment should lead to a greater influence of treatment expectation relative to external treatment^16^. From a statistical perspective, this would manifest as an interaction between agency and expectation, i.e. larger differences between low and high treatment expectations in self-treatment as compared to external treatment (see Fig. 1e).

Our data show clear effects of sensory attenuation and treatment expectations in two experiments with different pain rating modalities. At the same time, however, our data gives no indication of a significant interaction between sensory attenuation and treatment expectation effects, clearly favoring the forward model of perception for self-initiated pain treatment. This is corroborated by Bayesian model comparison, which strongly favored the forward model over the active inference model.

The Bayesian model representing the forward model included a free parameter (p_shift_) able to shift the mean of the likelihood distribution. One might argue that the likelihood, representing nociception, is a pure sensory signal which should be immune to any contextual modulation. However, given that the active inference model explicitly varies a term of the likelihood (i.e. precision), we decided to formulate the forward model in analogy, by also varying a likelihood component (i.e. mean). However, it should be noted that the forward model can be implemented in an alternative manner, namely using a free parameter that can vary the mean of the prior distributions instead of the mean of the likelihood.

To unravel neural mechanisms associated with top-down mechanisms of expectations and agency, EEG was measured during the second experiment. Here, we focused our analysis on two phases. Firstly, we investigated EEG power shortly after cue presentation, and secondly, we explored a time frame including the pain relief phase and the outcome phase, i.e. 1s before the target outcome temperature was reached by the thermode and 2s after.

Our analysis revealed a clear difference between VAS10 and VAS50 conditioning trials at the treatment outcome. Relative to the VAS50 condition, higher treatment success (VAS10) was associated with decreased alpha-to-beta (8-30Hz) and theta (4-7Hz) activity during the relief phase, while the outcome phase was associated with increased alpha-to-beta and decreased theta activity for VAS10 as compared to VAS50 trials. Interestingly, the same pattern emerged for self- versus external treatment trials, suggesting a similar mechanism for the sensory attenuation effect. Even though behavioral ratings indicated a comparable influence of treatment expectations, we did not find expectations associated with alterations of EEG activity at the treatment outcome. Instead, theta and alpha oscillations at the cue onset were differentially representing agency and expectations.

Oscillatory EEG responses after treatment do not represent a reliable correlate of pain – if this was the case, one would expect shared oscillatory mechanisms of all three pain suppressing behavioral effects (i.e. sensory attenuation, expectation and stimulus intensity). If there was a cortical oscillatory “pain matrix”^46–48^, both expectations and agency, which evidently modulate pain perception, should also modulate outcome-associated EEG responses. Here, we found modulations by stimulus intensity and agency, but not by expectations in outcome-locked EEG data. This is in line with previous studies which revealed cue-related expectation effects in the alpha-to-beta band before painful stimulation^44,45^, but not during painful stimulation. In another study, pain-induced alpha and gamma responses were significantly influenced by stimulus intensity but not by placebo hypoalgesia^49^. However, it has been demonstrated that expectation-based pain modulation can influence event-related pain potentials^43,49–52^.

Interestingly, oscillatory activity induced by agency (representing the sensory attenuation phenomenon) was comparable to differences in oscillatory patterns by different stimulus intensities (VAS10 versus VAS50) in conditioning trials, i.e. the same pattern emerged in different sets of trials. Here, sensory attenuation shares mechanisms with stimulus intensity processing, whereas expectation effects were not evident at the treatment outcome, but only after cue onset, which we attribute to cue-related processes. Following this line of thought, sensory attenuation might be directly related to alterations of oscillatory activity at pain-encoding areas associated with sensory processing. Expectation effects, on the other hand, might be encoded in oscillatory processes of brain areas typically associated with contextual influences of top-down processing.

In conclusion, pain treatment is additively enhanced by agency and positive expectations. Sensory attenuation and objectively different stimulus intensities both modify oscillatory activity at the relief and outcome phase of pain treatment, whereas expectation effects were associated with EEG activity directly following the cue. Using Bayesian model comparisons our data revealed no evidence for a decrease of precision in self-treatment, thus favoring a forward model as the mechanism underlying the positive effect of self-treatment.

## Methods

We conducted two experiments in which positive and negative treatment expectations as well as self- and external treatment were combined. In experiment 1, subjects were continuously rating their pain experience during painful stimulation and after self- or external treatment of pain. In experiment 2 with EEG recordings, we restricted the paradigm to include only two rating phases instead of a continuous rating to avoid excessive movement. Experiment 2 was preregistered at the German Clinical Trial Register (ID: DRKS00025541).

### Subjects

In experiment 1, 29 healthy participants were enrolled. All participants gave informed consent and were paid as compensation for their participation. Applicants were excluded if one of the following exclusion criteria applied: neurological, psychiatric, dermatological diseases, pain conditions, current medication, or substance abuse. All volunteers gave their informed consent. The study was approved by the Ethics Board of the Hamburg Medical Association. Four participants had to be excluded due to adverse reactions to the capsaicin cream, leaving a final sample of 25 participants (mean age 29.3, range 19–61 years, sex: 14 female / 11 male).

The required sample size of experiment 2 was determined according to a power calculation (G*Power V 3.1.9.4) based on the behavioral sensory attenuation and expectation effects in experiment 1. For the sensory attenuation effect, we observed an effect size of Cohen’s *f* = 0.508 (η_p_^2^ = 0.205) and an effect size of *f* = 0.669 (η_p_^2^ = 0.309) for the expectation effect. Using a power of (1-beta) of 0.8 and an alpha level of 0.05 and assuming low correlations (0.2) among repeated measures, this leads to a required sample size of 15, taking into account the weaker agency effect. However, given the different rating in the second experiment we increased the planned number of participants to 60. Assuming the same proportion of excluded participants as in experiment 1, this allowed us to potentially detect a medium effect size of *f* = 0.25^53^ with a sample size of 53.

We enrolled 60 healthy participants in experiment 2. Five participants had to be excluded due to adverse reactions to the capsaicin cream and 1 participant had to be excluded due to technical errors during recording, leaving a final sample size of 54 (mean 28.2, range 20–60 years, sex: 34 female / 20 male).

### Thermal stimulation and capsaicin application

Both experiments started with the same preparation procedure with the application of a capsaicin cream (ABC Heat Cream, Beiersdorf AG, 750µg capsaicin/g) to the left radial forearm. Two skin patches of the size of the thermal stimulator probe were covered with the capsaicin cream for a total of 15 minutes. Thermal stimulation was performed using a 30 × 30 mm^2^ Peltier thermode (Pathway model ATS, Medoc). The baseline temperature was set to 30°C for experiment 1 and the rise rate was set at 8°C/s for both experiments. The baseline temperature for experiment 2 was set at a lower temperature of 28°C to minimize skin irritation and attrition. After 2 blocks (experiment 1) or after the first experimental block (experiment 2), the capsaicin cream was reapplied for 5 minutes, and the stimulated skin patch was changed to avoid sensitization. In a first step, a single thermal stimulation with a slowly increasing ramp was used to familiarize the participant with the thermal stimulation. To test if the capsaicin cream was effectively reducing the pain threshold participants were asked to report the moment they felt a sensation of pain. If participants reported pain only above 46°C, the cream was reapplied for another 5 minutes on the skin patch (this applied to 2 participants in experiment 1 and to 4 participants in experiment 2).

### TENS illusion as a treatment situation

Afterwards, TENS (Transcutaneous Electric Nerve Stimulation) was established as a cover story for the treatment situation. TENS was presented as a nerve stimulation to effectively reduce pain by modulation of the nerve transmission. Putative TENS has been used to reliably generate treatment expectations in pain paradigms^19,54–56^. We provided volunteers with a deceptive “sham brochure” explaining that different stimulation frequencies result in different treatment efficacies. An electrode was attached to the elbow which was connected to an electrical current stimulator (Digitimer Ltd., model DS7A). Participants were told that the electrical current stimulator needed to be individually calibrated. For this, we applied short trains of electrical currents with increasing intensity and asked the participant to report if there was a sensation; this was intended to establish the belief that the device is actually active and capable of producing said currents. Afterwards and without knowledge of the participant, the electrical current stimulator was turned off and the participants were told that the settings for optimal stimulation were found. During the experiment, no actual electrical stimulation was applied. Additionally and to reinforce the TENS cover story, participants were asked to report if they felt a stimulation during the experiment and they were told that if this was the case, the stimulator needed to be recalibrated.

### Pain calibration

We individually calibrated the heat stimulation using an adaptive procedure to the levels of 10, 30, 50 and 70 on a Visual Analog Scale (VAS) from -100 to 100 where a VAS of 0 represented the pain threshold. The VAS was presented on a computer screen and ratings were given using the cursor keys on a conventional keyboard. At first, participants were stimulated with 34°C, 34.5°C, 35°C and 35.5°C and were asked to report if any of these stimuli were painful. Note that temperatures required to generate pain on capsaicin sensitized skin are in this temperature range. If the participant reported that the stimulation was painful, the procedure was continued with a starting temperature of 35°C, otherwise the starting temperature was set to 36°C for a stepwise procedure to find the pain threshold. Stimulus duration was set to 8s, according to the duration of VAS70 pain during the experiment. For the stepwise stimulus determination, 8 stimuli were presented with fixed reductions and increases in temperature relative to the pain threshold and participants were asked to rate the stimuli on a scale which was labeled as “normal sensation” at VAS -100, “minimally painful” at VAS 0 and “extremely painful” at VAS 100. Individual VAS levels of 10, 30, 50 and 70 were estimated using a linear regression of the VAS ratings recorded during this calibration phase. See Supplementary Information for calibration data.

### Trial design and block structure: experiment 1

Each trial was structured in 3 phases: Trial start, pain phase and treatment phase (see Fig. 2b). At trial start, an empty bar was presented in the center of the screen. The thermode temperature remained at the baseline of 30°C for 5s. Afterwards, the pain phase started, which was signaled by a filled red bar in the center of the screen. Thermode temperature was increased with a rate of 8°C/s to the temperature corresponding to the calibrated pain value of VAS70. The pain phase lasted for randomly jittered 8-10s. At the beginning of the treatment phase, a cue was presented which indicated whether a strong or weak treatment was to be expected and whether self-treatment or external treatment would occur. The cue was designed as a reduction of the centered red bar (i.e. more reduction to 1/3 of the total height with high treatment expectation as compared to 2/3 of the total height with low treatment expectation) and the word YOU (i.e. self-treatment) or HE (i.e. experimenter-induced, external treatment) written inside the bar, indicating self-or external treatment. After a lag of 2s, two treatment buttons were activated and appeared on the display, changing to green when pressed either by the subject or automatically. The external treatment was communicated as being done by the experimenter to reinforce the notion of a treatment setting, but was in fact computer initiated with a naturalistic “reaction time” delay. In the case of self-treatment, the participant pressed a button (A) and the treatment started with a reduction of the thermode temperature to the target level of VAS50, VAS30 or VAS10, depending on the condition. Meanwhile, participants received the signal of a button press (B) from the experimenter as an indication that the experimenter had acknowledged the self-treatment. In the case of an external treatment, a button press (B) by the participant had to acknowledge the external treatment. Meanwhile, experimental participants received an indication of a button press (A) from the experimenter, signaling that the treatment has been started. Participants were instructed to perform this task as soon as the treatment buttons appeared on the screen. Importantly, this procedure ensured identical motor output for the self-treatment and the external treatment conditions. In conditioning trials, the expectation of highly effective treatment resulted in a relatively more effective treatment and a reduction of the pain stimulus to the individual level of VAS10, as compared to a reduction of the pain stimulus to VAS50 in conditioning trials with low treatment expectations. For test trials, regardless of cued treatment effectivity, the pain stimulus was reduced to VAS30. The reduction of the pain stimulus was set at -8°C/s. In total, the treatment phase lasted 17-19s for a total trial duration of 32s including all three phases.

Additionally, participants were asked to continuously rate their pain level on a scale from 0 (minimal pain) to 100 (extreme pain) during the whole trial duration. A rating scale with a starting point at VAS0 (i.e. position of a red rating indicator) appeared ranging from VAS0, labelled as “minimally painful” to VAS100, labelled as “extremely painful”. The two buttons used for the rating were represented on the screen and were lighting up when pressed on the keyboard. After completion of the rating phase, the heat stimulus was reduced to the baseline temperature for the remaining intertrial interval of 18s.

During experiment 1, 4 experimental blocks were presented. Each block consisted of a total of 26 trials. It started with 8 conditioning trials, 4 of which were associated with low treatment success, 4 with high treatment success, each with the respective cues. After the conditioning trials, 3 micro blocks were presented consecutively. Each microblock consisted of 6 trials of following types:

(1) 1 conditioning reinforcement trial with high expectation of treatment success with actual high treatment success (reduction of pain from 70 to 10).
(2) 1 conditioning reinforcement trial as a reinforcement with low expectation to treatment success with actual low treatment success (reduction of pain from 70 to 50).
(3) 4 test trials with medium treatment success (reduction in pain from 70 to 30). I.e.:

a. 1 trial with high expectation of treatment success and self-treatment with a reduction of pain from 70 to 30.
b. 1 trial with high expectation of treatment success and external treatment with a reduction of pain from 70 to 30.
c. 1 trial with low expectation of treatment success and self-treatment with a reduction of pain from 70 to 30.
d. 1 trial with high expectation of treatment success and an external treatment with a reduction of pain from 70 to 30.

The order of trials within these micro blocks was randomized. Randomization was constrained so that a trial was not directly followed by the same type of trial, e.g. there were no two consecutive low expectation self-treatment test trials. For the 8 conditioning trials at the beginning of a block, it was ensured that at most two consecutive conditioning trials with the same condition (e.g. high treatment success) occurred.

In total, four experimental blocks were presented. During the first block conditioning and reinforcement trials were either exclusively self-treatment trials or external treatment trials. This was switched after two blocks, i.e. if the first two blocks were self-conditioning blocks the last two blocks were external conditioning blocks and vice versa.

Before the first experimental block was presented, 4 training trials were performed, during which the illusion of treatment was demonstrated by pressing the button connected to the heat stimulation device for pain reduction. At this stage, an actual button press by the experimenter was required during external training trials to establish the illusion of a direct link between TENS and the button press as for the participant. To do so, the experimenter sat next to the participant and demonstratively pressed the required button on the keyboard of the participant. In the remainder of the experiment, external “button presses” were performed by the computer unbeknownst to the subject.

### Trial design and block structure: experiment 2

During experiment 2 (see Fig. 2c), the paradigm was split into 5 phases: Trial start (4s), pain phase (8s), pain rating phase (6s), treatment phase (8s) and treatment rating phase (6s). Rating scales and related rating buttons on the screen were only presented during rating phases. During the pain rating phase, a red indicator on the VAS was presented with a random starting position. During the treatment rating phase, the final pain rating position of that red indicator was presented for orientation alongside a new green indicator initially appearing at a random position. The green indicator was used to rate the treatment outcome. In the treatment phase of experiment 2, treatment buttons were presented and activated simultaneously with the treatment cue without a jittered lag (as compared to experiment 1).

In total, 2 experimental blocks were presented. The first block consisted of conditioning and reinforcement trials which were either exclusively self-treatment trials or external treatment trials. This was switched after one block, i.e. if the first block was a self-conditioning block the second blocks was an external conditioning block, and vice versa. In total, 56 trials were presented per block, consisting of 8 conditioning trials followed by 8 micro blocks each containing both reinforcers (VAS10 and VAS50 conditioning trials) and each of the 4 test trial types (self- versus external conditioning, low versus high treatment expectation). Trials were presented with an intertrial interval of 4s. The first experimental block was presented after 4 training trials which were performed as in experiment 1.

### Questionnaire Data

After experiment 2 was concluded, participants were asked to complete several questionnaires. We included the BDI-V (Simplification of the Beck Depression Inventory)^57,58^, LSHS-R (Launay-Slade Hallucination Scale – German revised version)^59–61^, STAI-X2 (Trait) and STAI-X1 (State) (State-Trait-Anxiety-Scale)^62^, FKK (German locus of control scale, Fragebogen zu Kompetenz-und Kontrollüberzeugungen)^63^, SWE (German self-efficacy scale; Skala zur Erfassung der Selbstwirksamkeitserwartungen)^64^ and PCS (Pain Catastrophizing Scale - German translation)^65,66^ scales. Pearson product-moment correlation coefficients were computed to assess the relationship between questionnaire data and agency and placebo benefits. Agency benefits were defined as the difference between post-treatment VAS ratings of self-treatment and external treatment conditions. Placebo benefits were defined as the difference between post-treatment VAS ratings of high treatment expectation and low treatment expectation conditions. See Supplementary Information for a summary of the correlational results of the questionnaire data.

### EEG recording

EEG data were acquired using a 64-channel Ag/AgCl active electrode system (ActiCap64; BrainProducts) placed according to the extended 10–20 system^67^. Sixty electrodes were used of the most central scalp positions. The EEG was sampled at 500 Hz, referenced at FCz, and grounded at Iz. For removal of ocular movement artifacts, horizontal and vertical bipolar electrooculogram (EOG) were recorded using the four remaining electrodes.

### EEG preprocessing

The data analysis was performed using the Fieldtrip toolbox for EEG/MEG analysis^68^. For preprocessing, data were epoched and time-locked to the onset of the cue signaling the start of the treatment phase. Each epoch was centered (subtraction of the temporal mean) and included a time range of 19s before and 9s after trigger onset (starting with the empty cue signaling the start of the trial up to the end of the treatment phase).

We employed a preprocessing approach by Hipp et al. (2011) by splitting the data into two band-pass filtered sub-sets from 4 to 34Hz for low frequencies and from 16 to 250Hz for high frequencies^69^. This enabled efficient separation of low- and high frequency artifacts in subsequent ICA analysis. EEG epochs were visually inspected, and trials contaminated by artifacts due to gross movements or large technical artifacts were removed. Trials contaminated by eye-blinks, muscle activity, technical artifacts or movements were corrected using an independent component analysis (ICA) algorithm^70,71^ after careful inspection of topographies, power spectra, relation of ICA time courses to the temporal structure of the experiment and time-frequency representations. Artifactual components were removed before the remaining components were back-projected and resulted in corrected data. Subsequently, the data were re-referenced to a common average of all EEG channels and the previous reference channel FCz was reused as a data channel. Finally, epochs were visually screened and trials with remaining artifacts were excluded from analysis.

Before time–frequency transformations for data analysis were performed on the cleaned datasets, the time axis of single trials was shifted to create cue-locked and outcome-locked data. For cue-locked data, we set the onset of the cue signaling to the start of the treatment phase as t = 0. Outcome-locked data takes individual differences in response time into account and sets t = 0 to the time point when the thermode reached the treatment target temperature (calibrated VAS10, VAS30 and VAS50 levels for low conditioning, test and high conditioning trials, respectively). Trials were excluded if this duration (from cue onset to treatment outcome) was longer than 6s. This allowed us to create an analysis window of 2s in subsequent time-frequency analysis without contamination by the subsequent rating phase.

### EEG spectral analysis

Spectral analysis was adapted from Hipp et al. (2011)^69^. This approach ensured a homogenous sampling and smoothing in time and frequency space. We calculated spectral estimates for 23 logarithmically scaled frequencies ranging from 4 to 181 Hz (0.25 octave increments) for the pain phase and treatment phase in 0.05s steps. For cue-locked data, this included the treatment phase from cue onset up to 2s after cue onset. For treatment-locked data, this included the relief phase from 1s before the treatment outcome (target temperatures of VAS10, VAS30 or VAS50) was reached, and the outcome phase up to 2s after the target temperature was reached. Using the multitaper (DPSS) approach, we set the temporal and spectral smoothing to match 250ms and 3/4 octave, respectively. For frequencies below 16 Hz, we employed 250ms temporal windows and varied the number of Slepian tapers to approximate a 3/4 octave spectrum smoothing. We changed the time window for frequencies below 16 Hz to achieve a frequency smoothing of 3/4 octaves with a single taper. We computed the frequency transform using high- and low-frequency data for frequencies above and below 25 Hz, respectively. Analysis was then continued with the combined spectral data after averaging of spectral estimates per block and condition over trials for each subject.

For the baseline correction of time–frequency data, the mean and standard deviation were estimated (for each subject/channel/frequency combination, separately) from 0.5 to 7.5s of the pain phase (i.e. increases and decreases in EEG power activity indicate deviations from EEG power during painful stimulation). The mean spectral estimate of the baseline was then subtracted from each data point, and the resulting baseline-centered values were divided by the baseline standard deviation (classical baseline normalization – additive model; see Grandchamp & Delorme, 2011)^72^.

### Behavioral data analysis

For experiment 1, we performed analysis on the continuous VAS rating by simply using the last data point of each trial. For experiment 2, we performed analysis on the single post-treatment VAS rating. Here, we have 2×2 conditions for test trials (low versus high treatment expectation / self- versus external treatment) and 2 conditions for conditioning trials. Firstly, we conducted a 3×1 repeated measures ANOVA with post-hoc t-tests to evaluate the differences between VAS10 (conditioning), VAS30 (test) and VAS50 (conditioning) conditions, respectively. Secondly, we conducted a 2×2 repeated measures ANOVA to evaluate differences between the different test conditions (low versus high expectation / self- versus external treatment) and the interaction term of expectation and agency.

### Bayesian integration models of placebo pain treatment

For model-based analysis of our post-treatment VAS rating data, we designed two Bayesian integration models of pain perception in placebo pain treatment (see Büchel et al., 2014 for a review)^16^ in accordance with the forward model and active inference model. In the Bayesian formulation of pain perception, Bayes’ theorem is used to estimate the level of perceived pain, taking precision-weighted prior experiences into account (Fig. 1a and Eq.1). Formally, the model integrates a prior with a likelihood to estimate a posterior. Both, the prior and the likelihood were approximated by normal distributions allowing for an analytical integration using normal-normal conjugate priors to estimate the normal posterior.

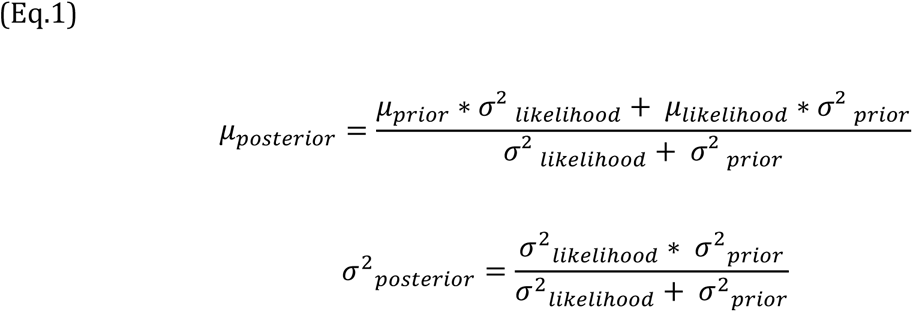

With respect to the behavioral data, our model predicted the painfulness of the test phase post-treatment VAS ratings (posterior) by integrating conditioning post-treatment VAS ratings as a prior (mean and variance derived from VAS10 and VAS50 post-treatment VAS ratings for high and low treatment expectation conditions, respectively) with an individual estimate of the likelihood (average of VAS10 and VAS50 parameters for each subject). Gaussian approximation of the rating data was performed by fitting a Gaussian cumulative probability density functions to the cumulative sum of the ratings using a robust grid search^19^.

For the estimation of the posterior parameters in self-treatment trials we created two derived models, based on predictions of the forward model and the active inference model. For the forward model, we included a free parameter *p_shift_* to enable a likelihood shift (Eq.2; Fig. 1b):

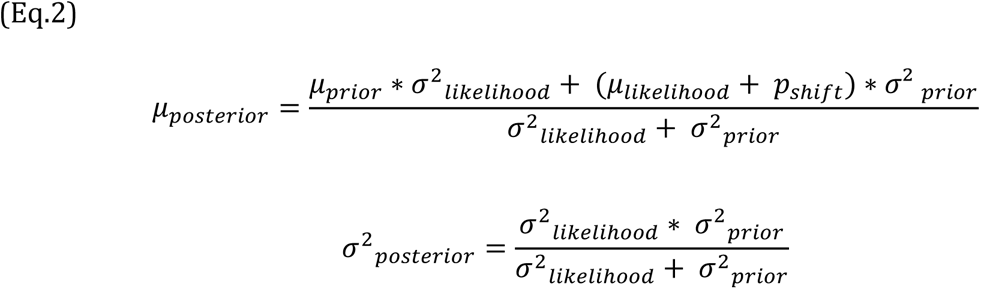

Active inference assumes a higher likelihood variance (i.e. lower precision) in self-treatment trials, and thus we included a free parameter *p_relax_* in the active inference model to represent a modulation of likelihood variance by self-treatment. Under active inference, posterior parameters are estimated by the following equations (see also Fig. 1c):

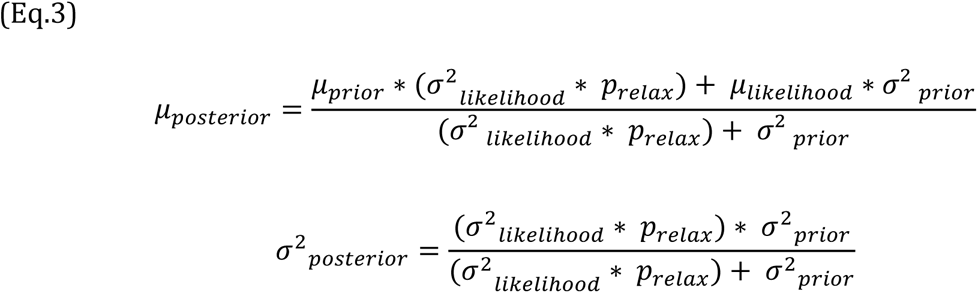

We used a variational Bayesian inference to estimate the parameters of all models using the VBA toolbox^73^ for Matlab (R2021a). We used uninformative priors for both parameters (p_relax_ ∼ *Normal*(1,1000) and p_shift_ ∼ *Normal*(0, 1000). In addition, we fitted a combined model with both free parameters and a null model (Eq.1) in which both parameters were “constrained” through their priors (p_relax_ ∼ *Normal*(1, 1e-20) and p_shift_ ∼ *Normal*(0, 1e-20). Given our behavioral post-treatment VAS rating data (i.e. empirical posterior), VBA recovers an approximation to both the posterior density on unknown variables (p_relax_ and p_shift_ for the active inference model and the forward model, respectively) and the log model evidence (which is used for model comparison). We used a random effects (RFX) Bayesian model selection approach^38,39^ to estimate the overall posterior model probability across subjects. Finally, we estimated the protected exceedance probability as a metric for the Bayesian model comparison of the active inference model and the forward model^38^.

### EEG data analysis

Here, we analyzed the effects of two phases of pain treatment. Firstly, we wanted to analyze effects associated with the treatment cue indicating low or high treatment success and self- or external treatment. For this analysis, we used a cue-locked analysis window of 2s after the onset of the cue. Secondly, we wanted to evaluate the relief phase and treatment outcome based on low or high treatment expectations, agency, and their interaction. As the treatment outcome occurred at highly variable time points based on the response, we analyzed –1 to 2s in relation to the time point when the thermode reached the treatment target temperature (where -1 to 0s was defined as the relief phase and 0 to 2s was defined as the treatment outcome phase).

We corrected all statistical tests in electrode space for multiple comparisons using non-parametrical permutation tests of clusters^74^. Samples (exceeding the threshold of *p* < .05) were clustered in connected sets on the basis of temporal (i.e. adjacent time points), spatial (i.e. neighboring electrodes), and spectral adjacency. Clustering was restricted in a way that only samples were included in a cluster which had at least one significant neighbor in electrode space (i.e. at least one neighboring channel also had to exceed the threshold for a sample to be included in the cluster). Neighbors were defined by a template provided by the Fieldtrip toolbox corresponding to the used EEG montage.

A cluster value was defined as the sum of all statistical values of included samples. Monte Carlo sampling was used to generate 1000 random permutations of the design matrix, and statistical tests were repeated in time–frequency space with the random design matrices. The probability of a cluster from the original design matrix (p-value) was calculated by the proportion of random design matrices producing a cluster with a cluster value exceeding the original cluster. Muscular and ocular electrodes were excluded from the cluster analysis.

Further, we wanted to explore correlations of between-subject time-frequency responses and benefits of high treatment expectation versus low treatment expectation in post-treatment VAS ratings, as well as sensory attenuation model parameters. For each participant, we calculated the placebo benefit by the within-subject z-normalized difference of behavioral ratings between high and low treatment expectations in test trials. For the benefit of agency, we used the single subject mean estimate of the p_shift_ parameter from the VBA model inversion procedure, accordingly.

Any positive or negative cluster in correlation analysis of EEG power during test trials would indicate an association with placebo or agency benefits. Here, a p-value of p < 0.05 obtained from the Pearson’s correlation test statistic as implemented in the Fieldtrip toolbox was used as a threshold for clustering.

## Acknowledgements

CB is supported by ERC-AdG-883892-PainPersist and DFG SFB 289 project A02. MR is supported by DFG SFB 289 project A03 and DFG SFB TR 169 project B3. Funded by the Deutsche Forschungsgemeinschaft (DFG, German Research Foundation) – Project-ID 422744262–TRR 289.

## Ethics

Human subjects: All volunteers gave their informed consent. The study was approved by the Ethics board of the Hamburg Medical Association (PV3892). The authors report no conflict of interest.

## Author contributions

A.S.: Conceptualization, Data curation, Software, Formal analysis, Investigation, Visualization, Methodology, Writing - original draft, Project administration, Writing - review and editing; B.H.: Conceptualization, Software, Methodology, Writing - review and editing; M.R.: Resources, Methodology; C.B.: Conceptualization, Resources, Formal analysis, Supervision, Funding acquisition, Validation, Visualization, Methodology, Project administration, Writing - review and editing.

## Data availability

Data for this study are available on https://osf.io/q8tgj/

Strube A., Horing B., Rose M., Büchel C., (2022) Open Science Framework ID q8tgj. Placebo and Sensory Attenuation in Pain Treatment.

## Supplementary Information

### Calibration data: experiment 1

During experiment 1, pain levels were calibrated to achieve VAS10 (M = 38.1°C, SD = 3.5°C, Min = 31.8°C, Max = 44.8°C), VAS30 (M = 39°C, SD = 3.5°C, Min = 32.2°C, Max = 45.3°C), VAS50 (M = 39.9°C, SD = 3.6°C, Min = 32.5°C, Max = 46.2°C) and VAS70 (M = 40.8°C, SD = 3.8°C, Min = 32.8°C, Max = 47.2°C) pain levels (for highly effective conditioning, test trials, low effective conditioning and VAS70 pain stimulation, respectively).

### Calibration data: experiment 2

During experiment 2, pain levels were calibrated to achieve VAS10 (M = 38.2°C, SD = 3.1°C, Min = 31.72°C, Max = 44.5°C), VAS30 (M = 39°C, SD = 3.1°C, Min = 32.1°C, Max = 45.3°C), VAS50 (M = 38.19°C, SD = 3.1°C, Min = 32.5°C, Max = 46.1°C) and VAS70 (M = 40.5°C, SD = 3.2°C, Min = 32.8°C, Max = 46.9°C) pain levels (for highly effective conditioning, test trials, low effective conditioning and pain stimulation, respectively).

### EEG data analysis: interaction

We conducted post-hoc t-tests to confirm the crossed interaction of agency and expectations at cue-locked EEG data. Post-hoc t-tests confirmed a crossed interaction where all 4 comparisons were significant, i.e. self-treatment with high treatment expectation trials were associated with larger EEG power than self-treatment trials with low treatment expectations (t(53) = 3.74, p < 0.001), whereas external treatment trials with high treatment expectations were associated with smaller EEG power than external treatment trials with low treatment expectations (t(53) = -5.17, p < 0.001). Also, high treatment expectation trials with self-treatment were associated with higher EEG power that high treatment expectation trials with external treatment (t(53) = 5.84, p < 0.001) and low treatment expectation trials with self-treatment were associated with lower EEG power than low treatment expectation trials with external treatment (t(53) = 3.04, p = 0.004).

### Questionnaire data

We correlated z-scored agency VAS benefits (self-treatment – external treatment post-treatment VAS rating) and placebo VAS benefits (high treatment expectation – low treatment expectation post-treatment VAS rating) with scores from various questionnaire data (BDI_V, FKK, LSHS_R, PCS, STAI-X2 (Trait), STAI-X1 (State), SWE). All correlations were not significant (all *p* > 0.05, uncorrected; Supplementary Table 1).

**Supplementary Table 1.**
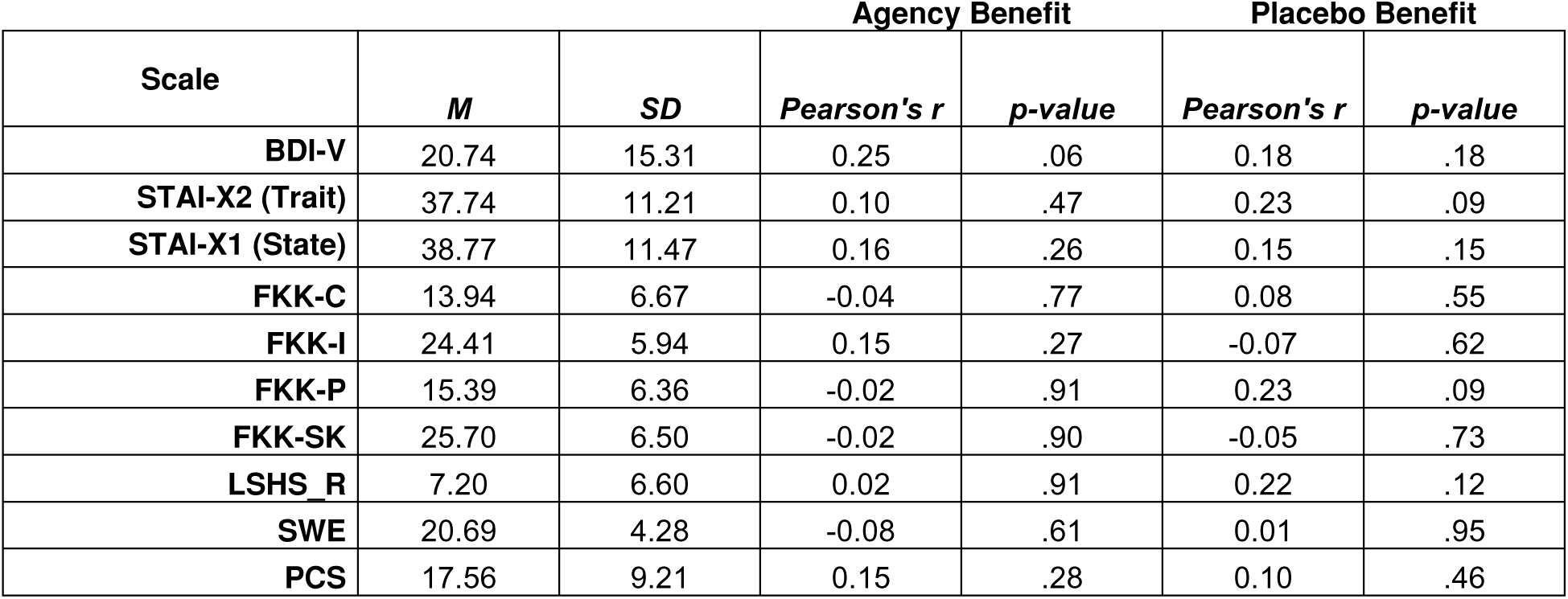
Correlation of questionnaire data with individual agency and placebo benefit scores.

## Supplementary Figures

**Supplementary Figure 1.**
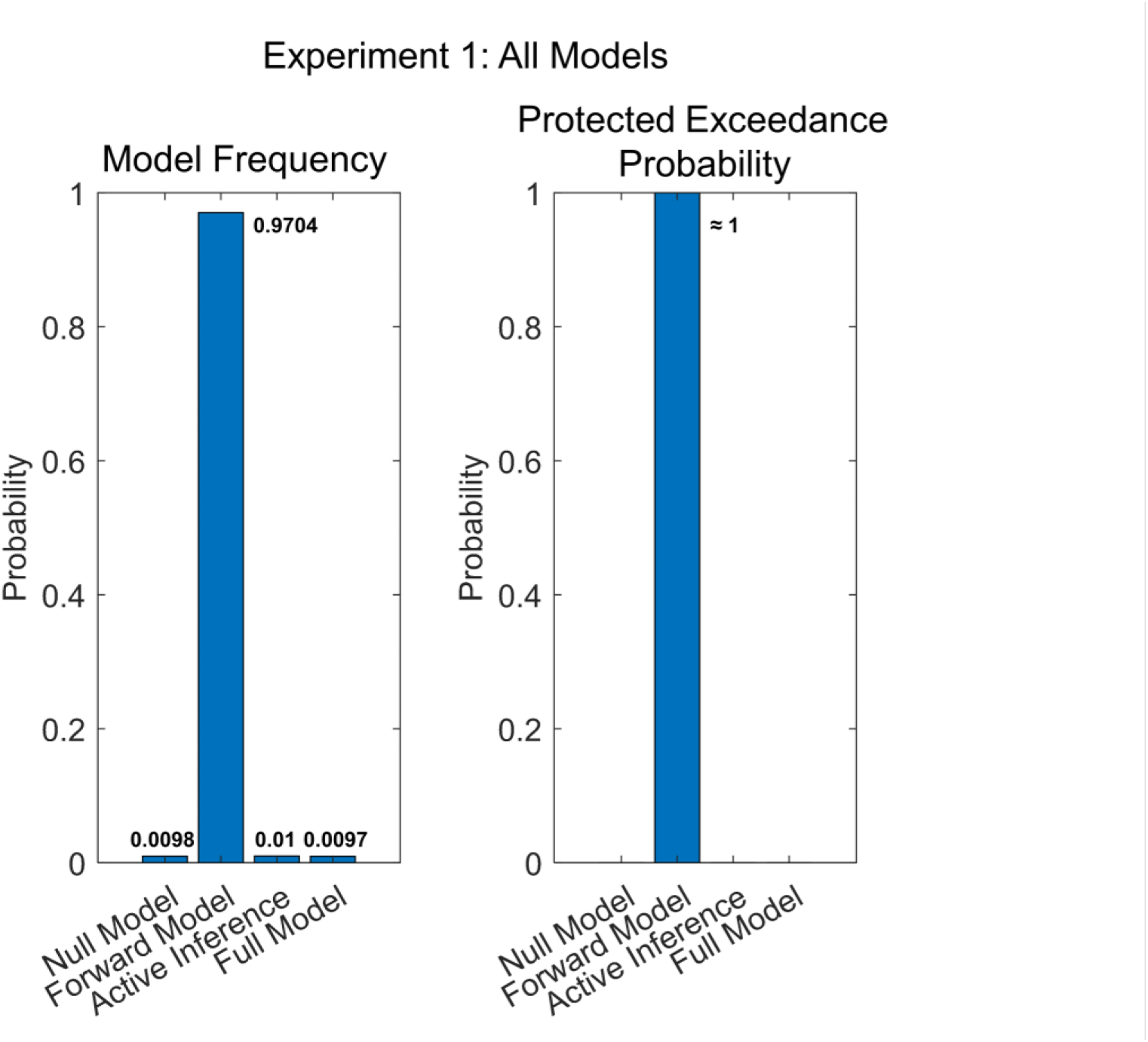
Results of the Bayesian model comparison including all models (null model, forward model, active inference and full model) of experiment 1 (N=25). Model frequencies (left) and protected exceedance probabilities (right).

**Supplementary Figure 2.**
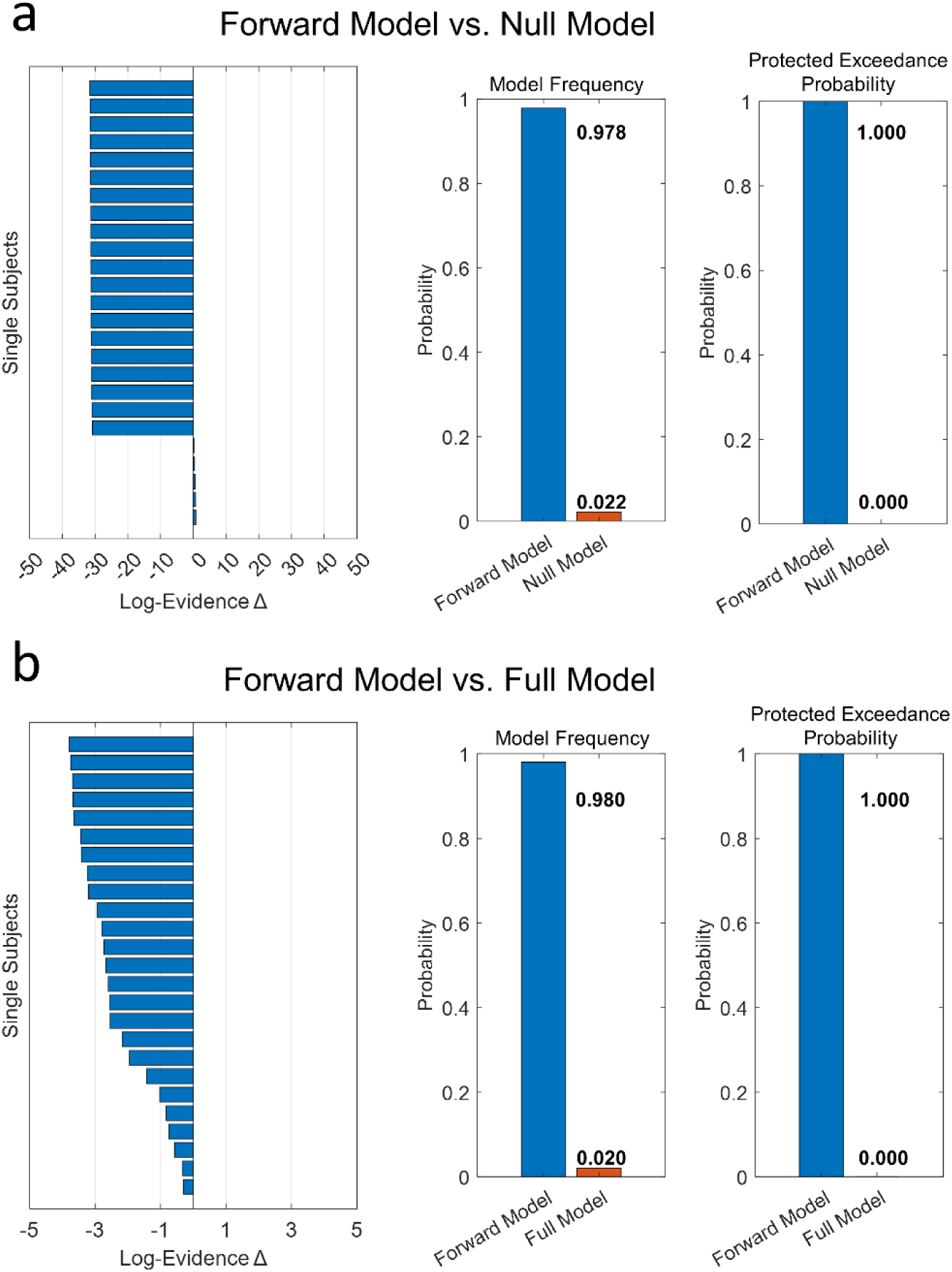
Results of the Bayesian model comparison of (a) forward model versus null model and (b) forward model versus full model of experiment 1 (N=25). Single subject differences of log evidence for the forward model versus null/full model (negative values favor the forward model) (left), model frequencies (center), and protected exceedance probabilities (right).

**Supplementary Figure 3.**
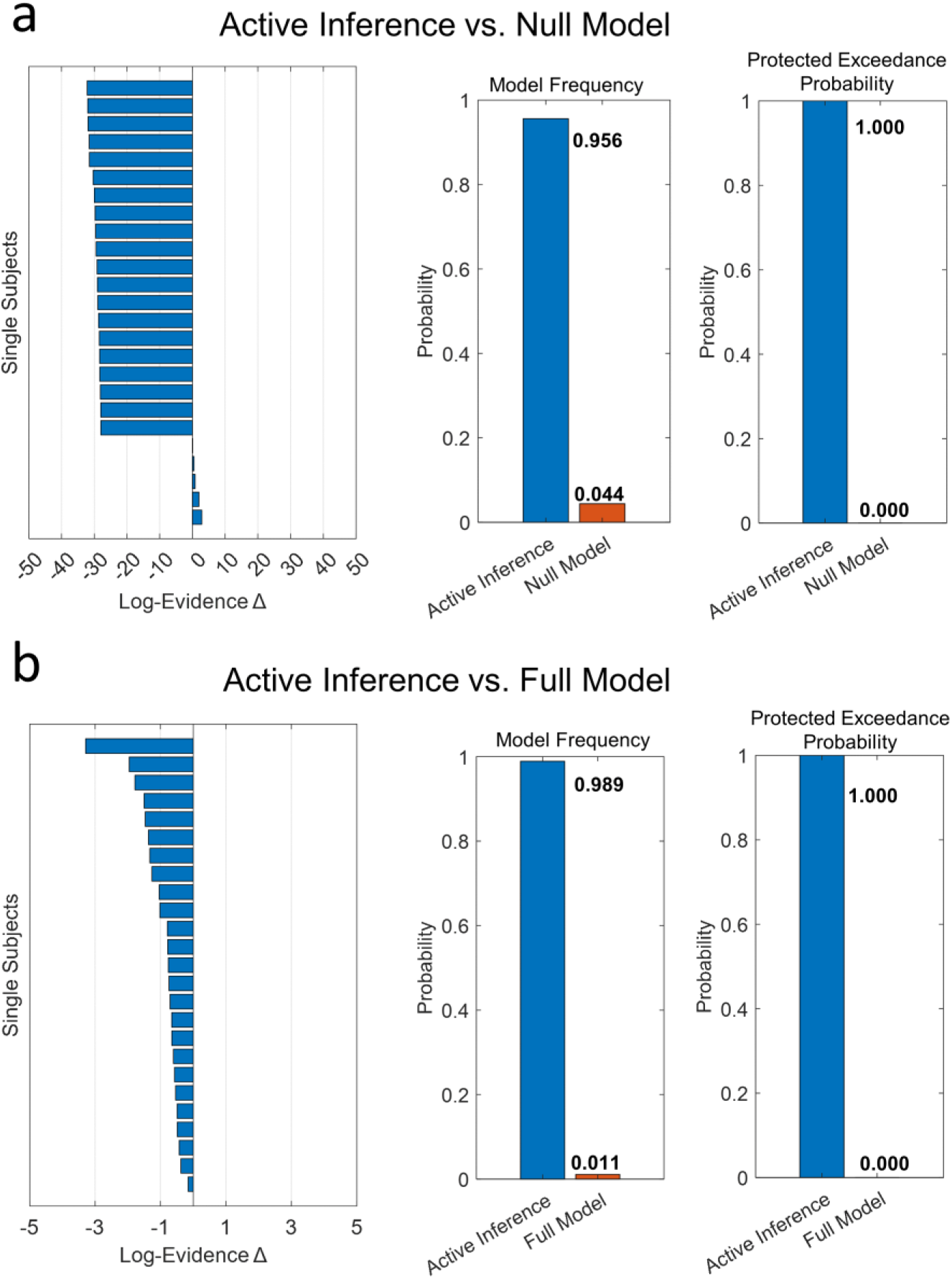
Results of the Bayesian model comparison of (a) active inference versus null model and (b) active inference versus full model of experiment 1 (N=25). Single subject differences of log evidence for the active inference versus null/full model (negative values favor the active inference model) (left), model frequencies (center), and protected exceedance probabilities (right).

**Supplementary Figure 4.**
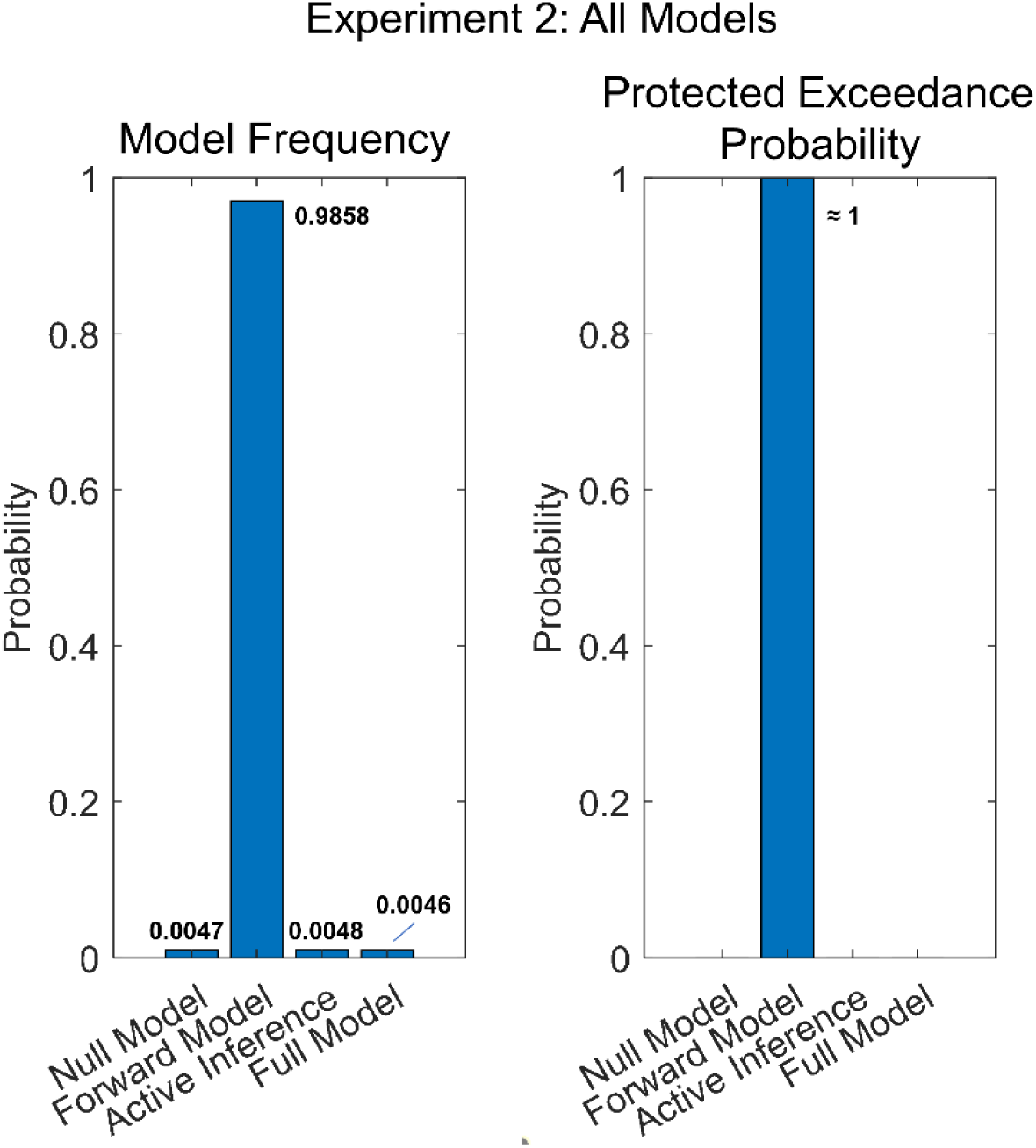
Results of the Bayesian model comparison including all models (null model, forward model, active inference and full model) of experiment 2 (N=54). Model frequencies (left) and protected exceedance probabilities (right).

**Supplementary Figure 5.**
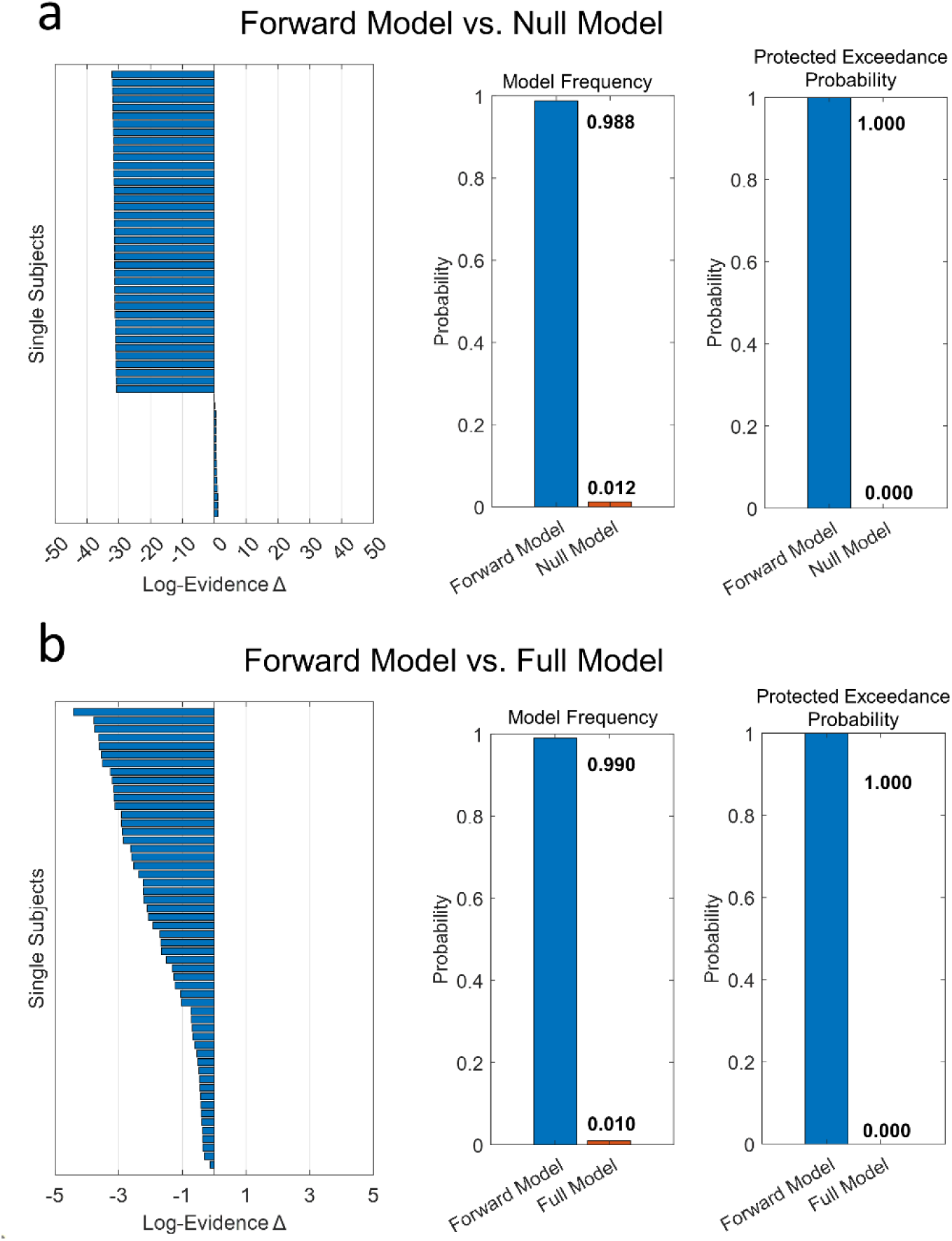
Results of the Bayesian model comparison of (a) forward model versus null model and (b) forward model versus full model of experiment 2 (N=54). Single subject differences of log evidence for the forward model versus null/full model (negative values favor the forward model) (left), model frequencies (center), and protected exceedance probabilities (right).

**Supplementary Figure 6.**
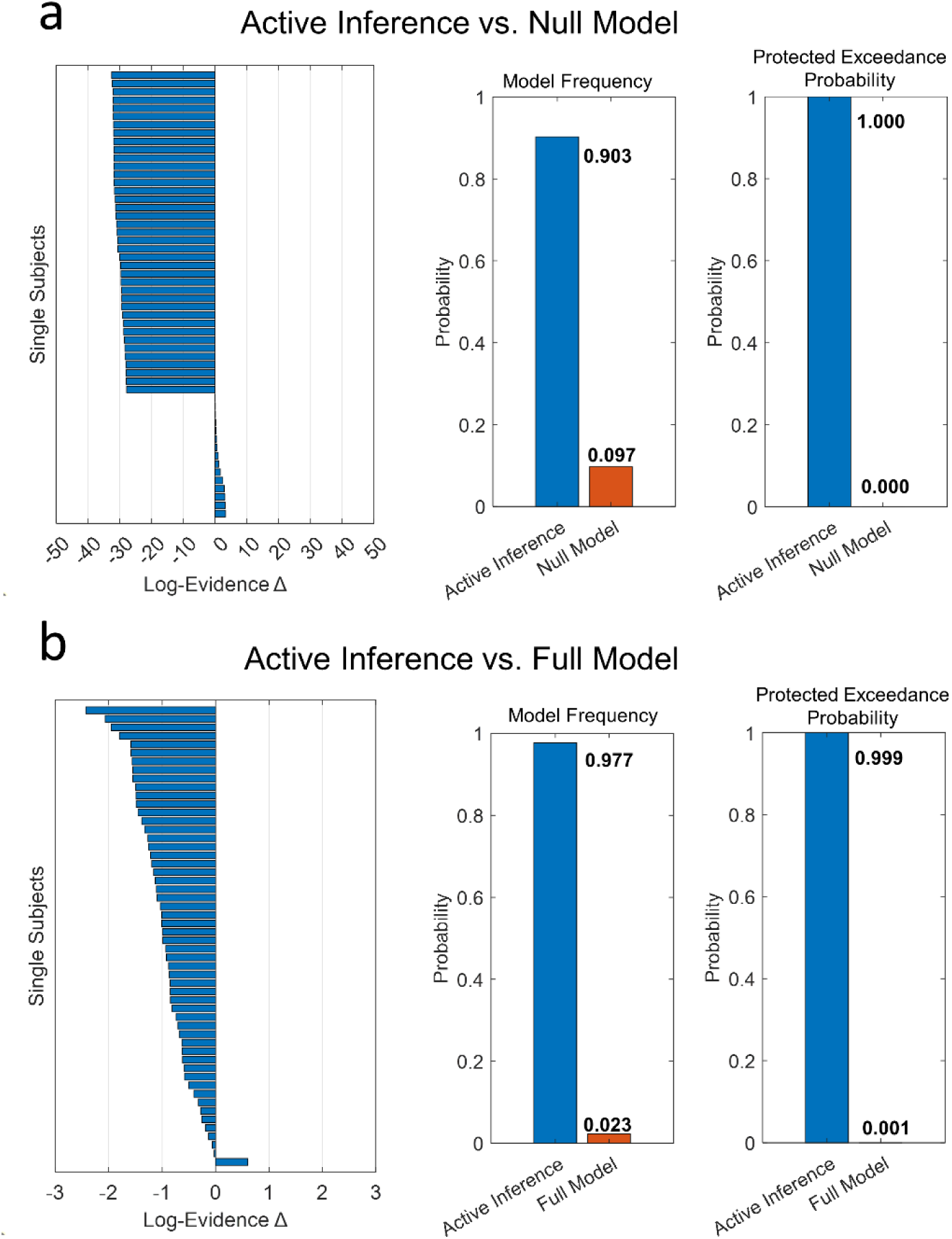
Results of the Bayesian model comparison of (a) active inference versus null model and (b) active inference versus full model of the experiment 2 (N=54). Single subject differences of log evidence for the active inference versus null/full model (negative values favor the active inference model) (left), model frequencies (center), protected exceedance probabilities (right).

**Supplementary Figure 7.**
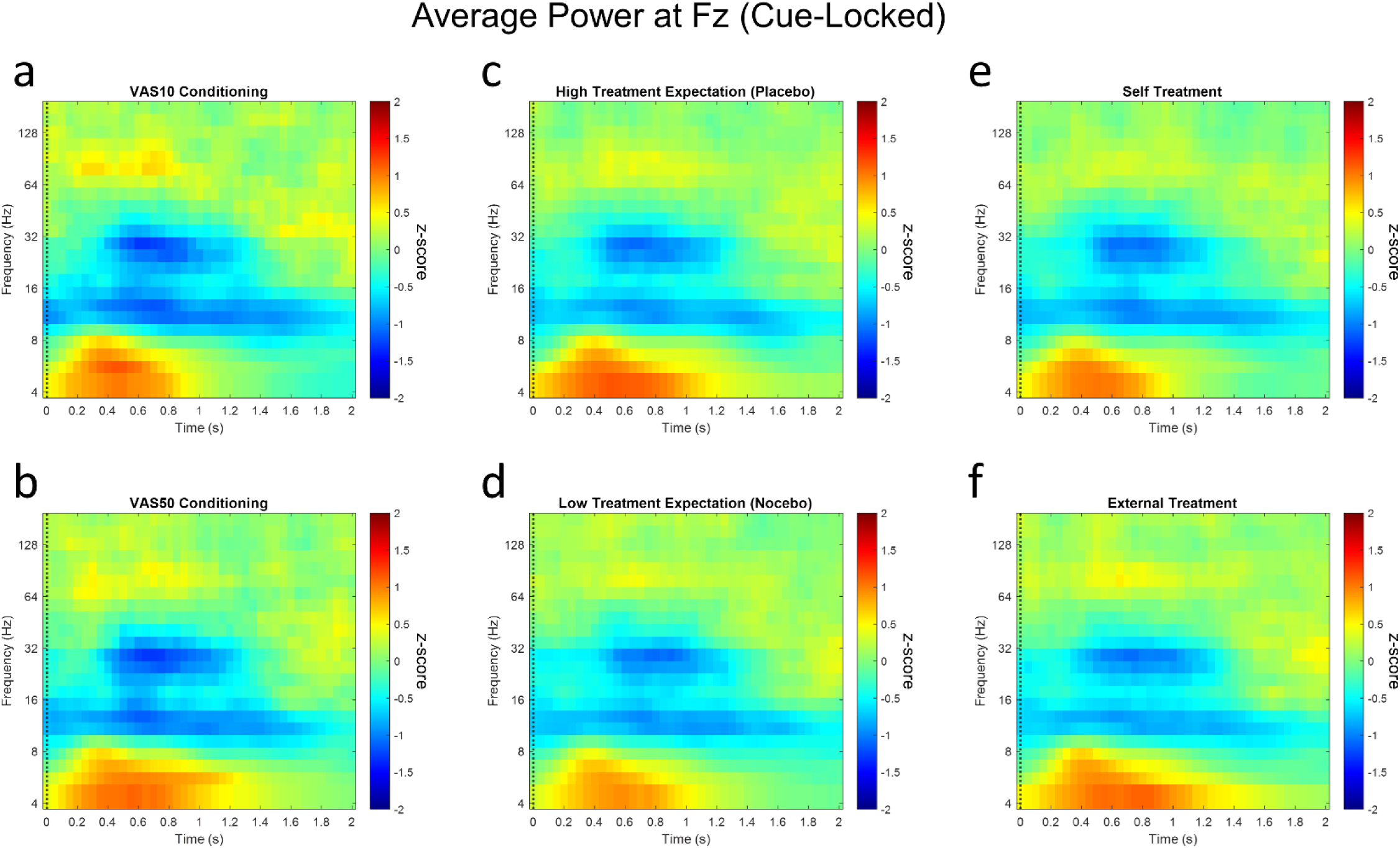
Time-frequency plots represent averaged, baseline-corrected, cue-locked power values for each condition at Fz. (a) VAS10 versus (b) VAS50 conditioning, (c) high versus (d) low treatment expectation and (e) self- versus (f) external treatment. Warm colors represent positive z-values (increase of power compared to baseline) and cold colors represent negative z-values (decrease of power compared to baseline).

**Supplementary Figure 8.**
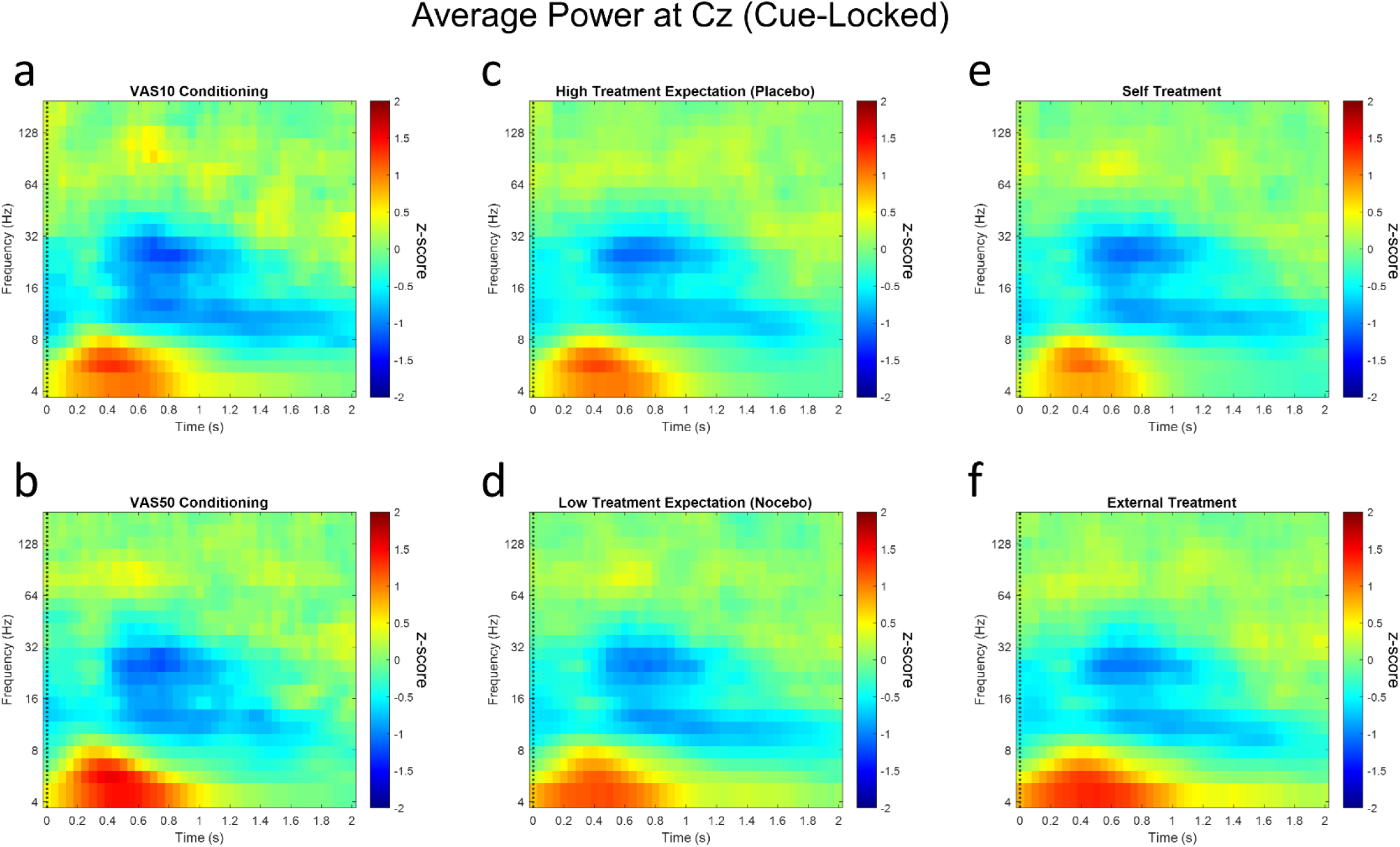
Time-frequency plots represent averaged, baseline-corrected, cue-locked power values for each condition at Cz. (a) VAS10 versus (b) VAS50 conditioning, (c) high versus (d) low treatment expectation and (e) self- versus (f) external treatment. Warm colors represent positive z-values (increase of power compared to baseline) and cold colors represent negative z-values (decrease of power compared to baseline).

**Supplementary Figure 9.**
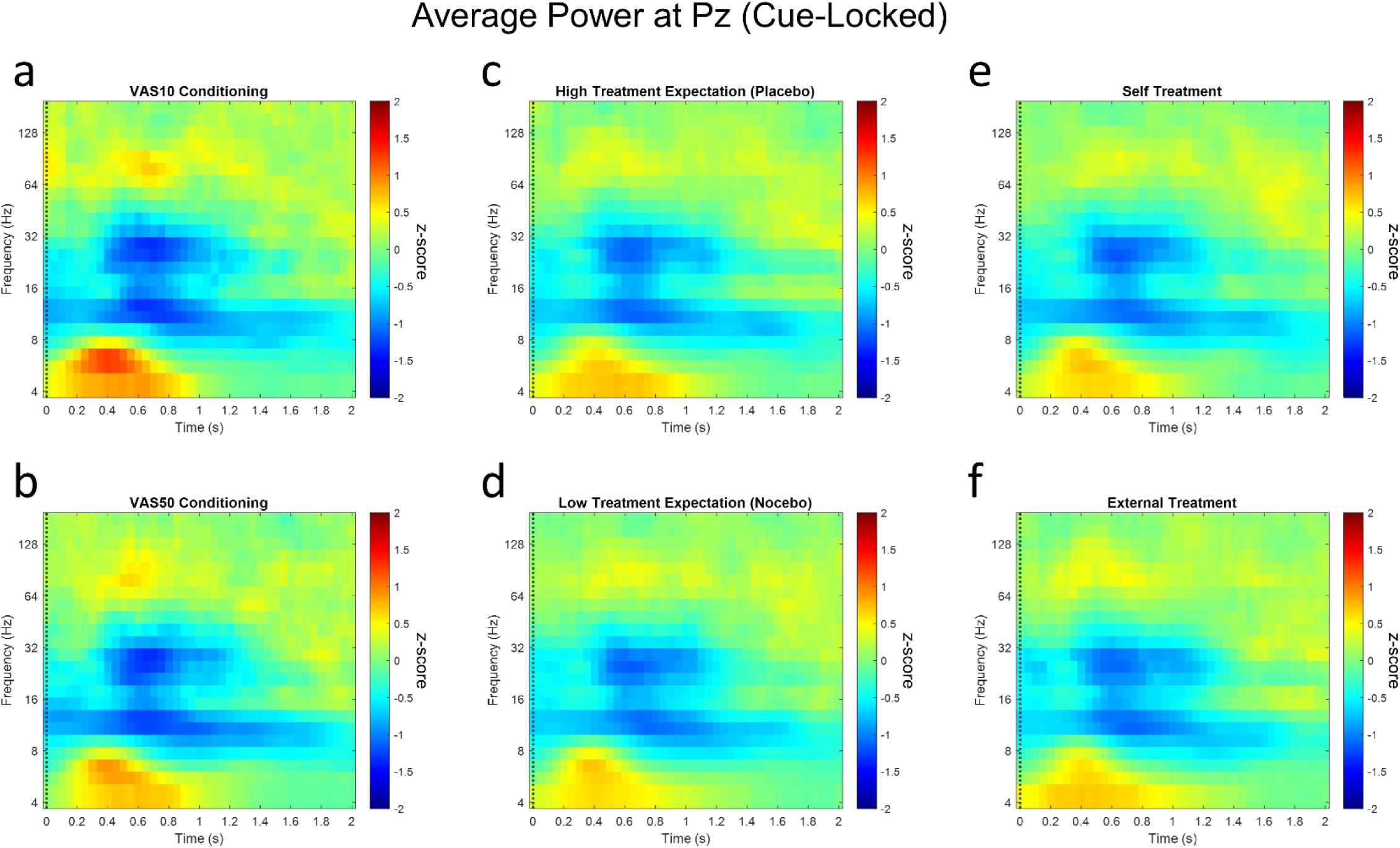
Time-frequency plots represent averaged, baseline-corrected, cue-locked power values for each condition at Pz. (a) VAS10 versus (b) VAS50 conditioning, (c) high versus (d) low treatment expectation and (e) self- versus (f) external treatment. Warm colors represent positive z-values (increase of power compared to baseline) and cold colors represent negative z-values (decrease of power compared to baseline).

**Supplementary Figure 10.**
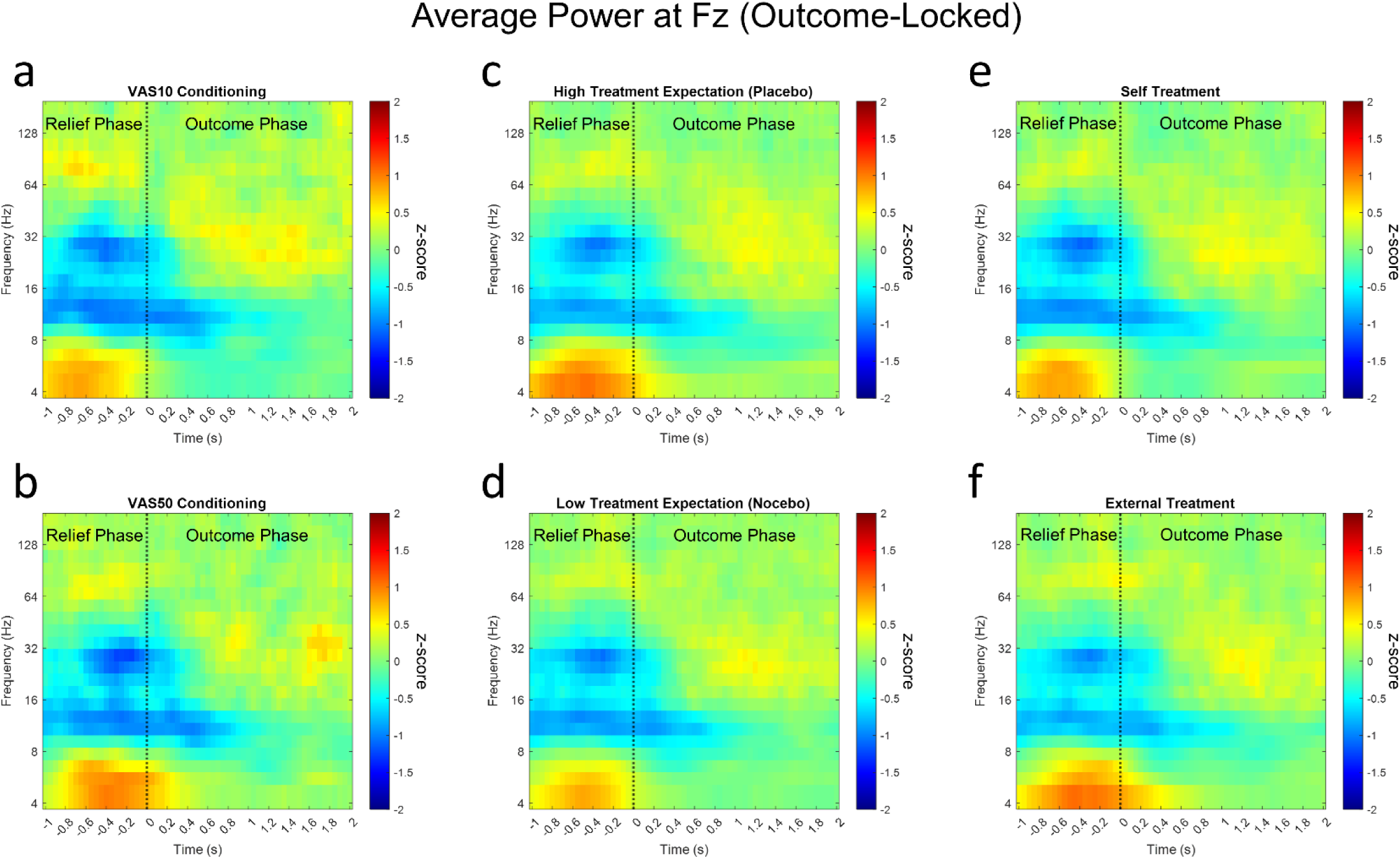
Time-frequency plots represent averaged, baseline-corrected, outcome-locked power values for each condition at Fz. (a) VAS10 versus (b) VAS50 conditioning, (c) high versus (d) low treatment expectation and (e) self- versus (f) external treatment. Warm colors represent positive z-values (increase of power compared to baseline) and cold colors represent negative z-values (decrease of power compared to baseline).

**Supplementary Figure 11.**
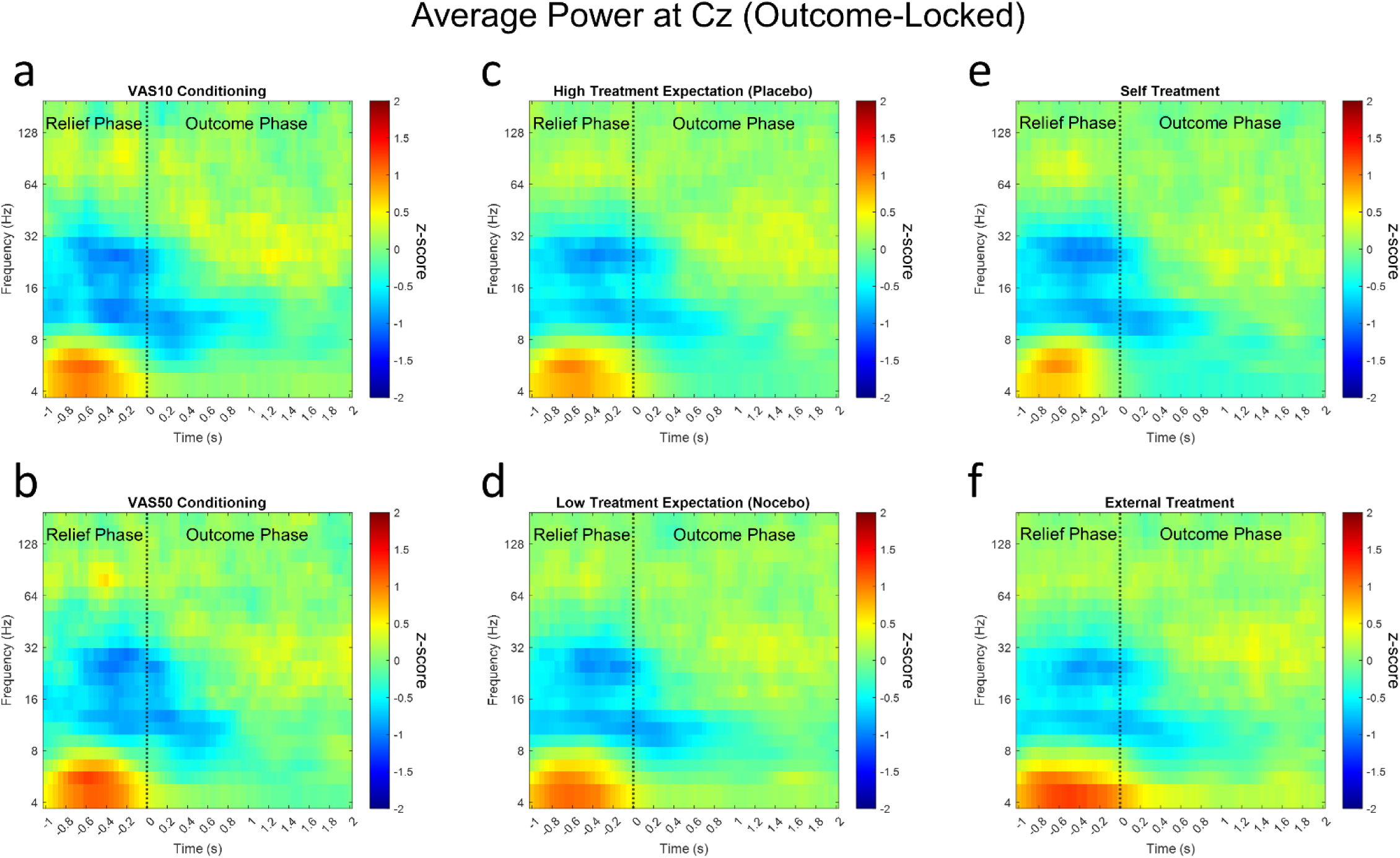
Time-frequency plots represent averaged, baseline-corrected, outcome-locked power values for each condition at Cz. (a) VAS10 versus (b) VAS50 conditioning, (c) high versus (d) low treatment expectation and (e) self- versus (f) external treatment. Warm colors represent positive z-values (increase of power compared to baseline) and cold colors represent negative z-values (decrease of power compared to baseline).

**Supplementary Figure 12.**
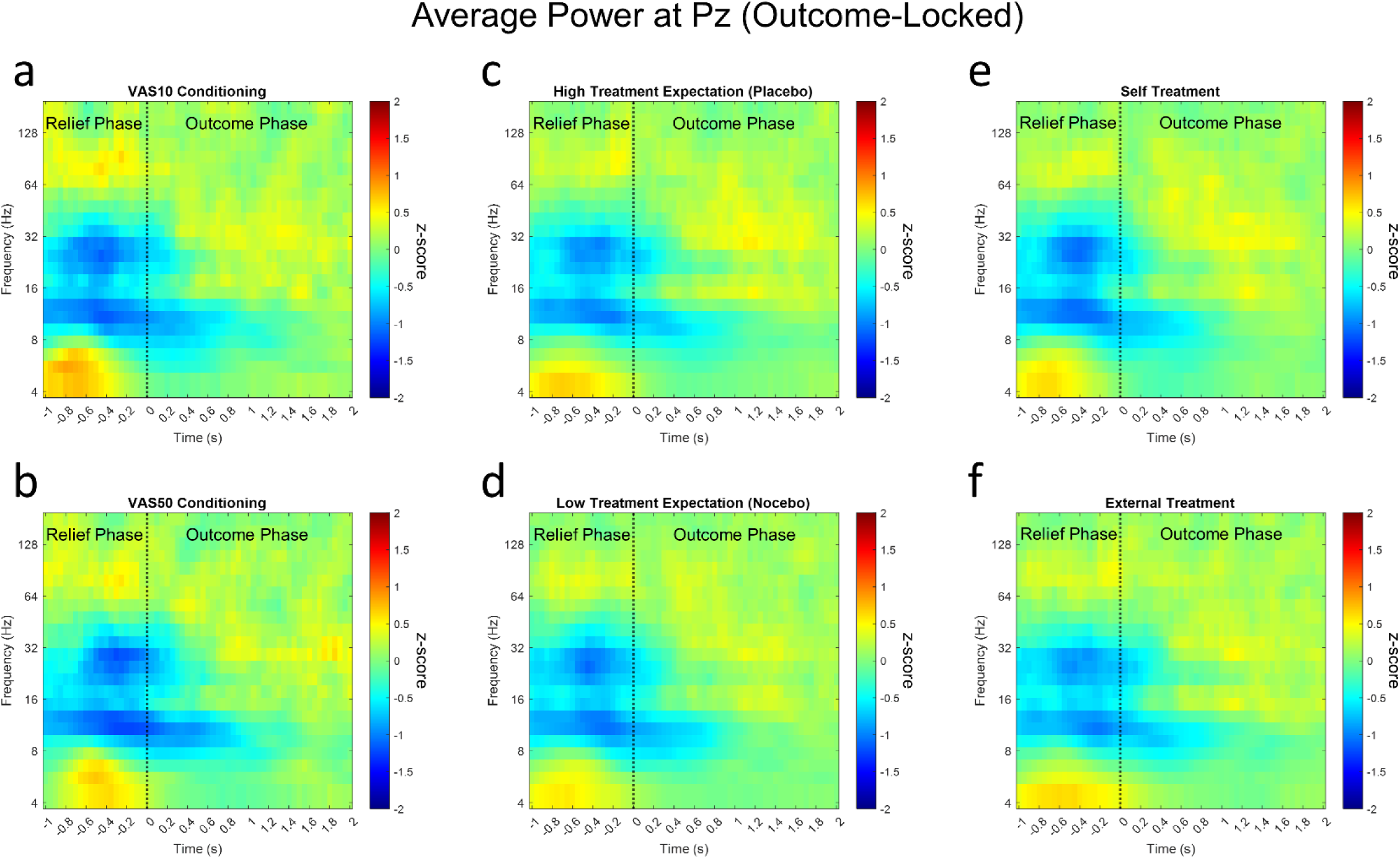
Time-frequency plots represent averaged, baseline-corrected, outcome-locked power values for each condition at Pz. (a) VAS10 versus (b) VAS50 conditioning, (c) high versus (d) low treatment expectation and (e) self- versus (f) external treatment. Warm colors represent positive z-values (increase of power compared to baseline) and cold colors represent negative z-values (decrease of power compared to baseline).

**Supplementary Figure 13.**
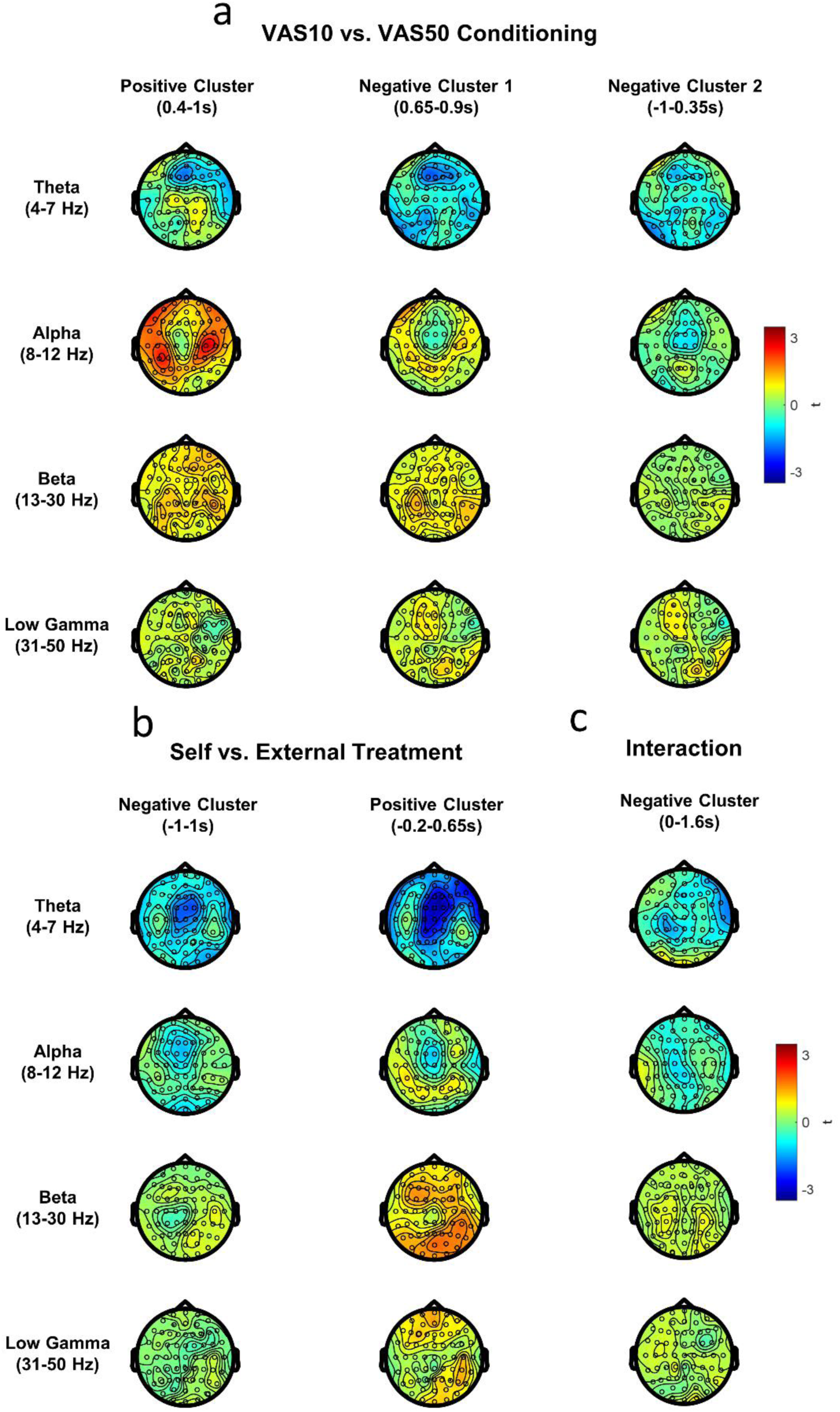
Topographies represent averaged t-values of pre-defined frequency bands (Theta 4-7Hz, Alpha 8-12Hz, Beta 13-30Hz and Low Gamma 31-50Hz) over the time range of significant clusters of (a) VAS10 versus VAS50 conditioning, (b) self- versus external treatment and (c) the interaction of agency and expectation.

**Supplementary Figure 14.**
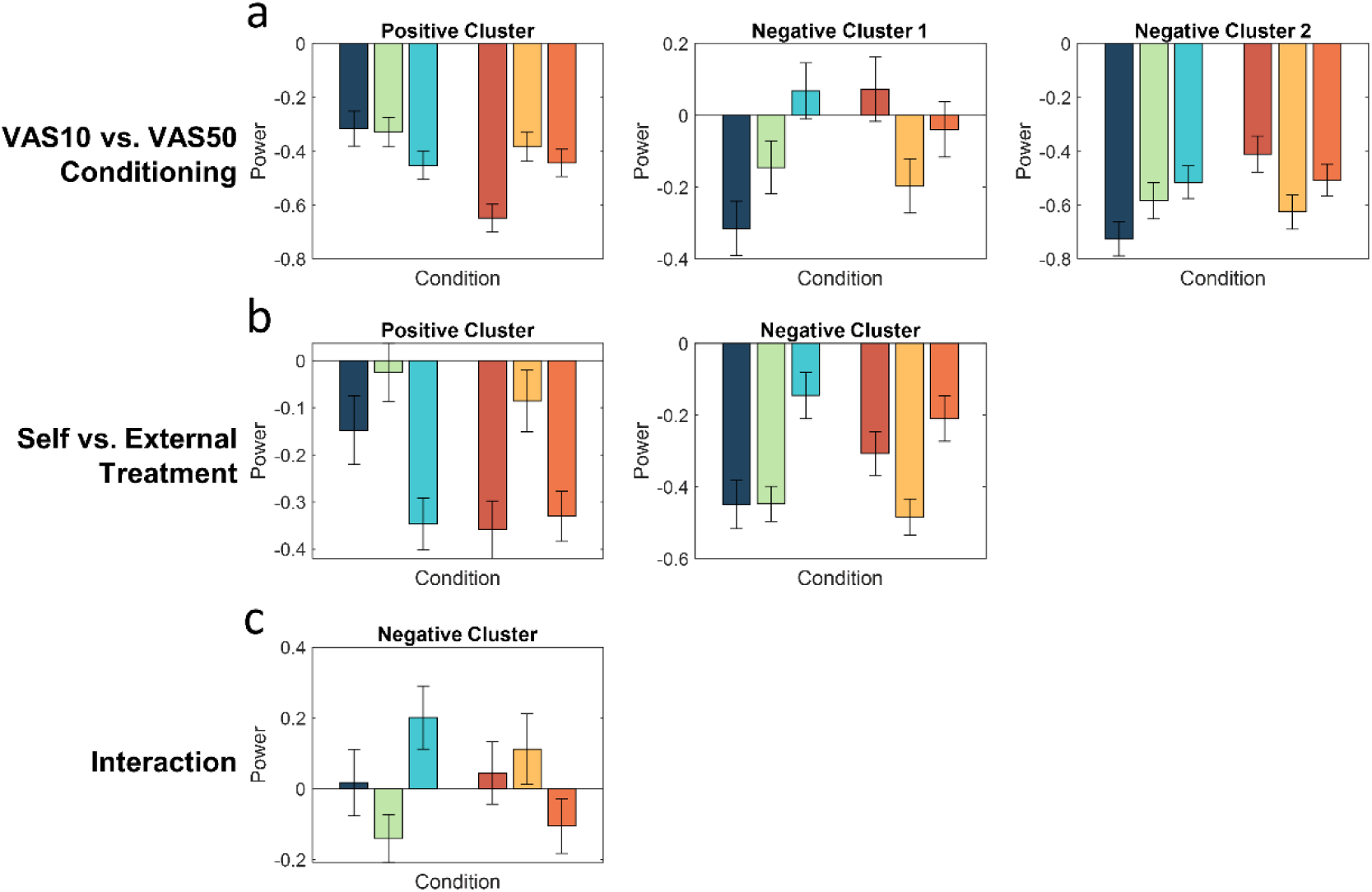
Bar graphs represent averaged (baseline-corrected) power values for each condition averaged over all data points included in significant clusters of (a) VAS10 versus VAS50 conditioning, (b) self- versus external treatment and (c) the interaction of agency and expectation. Error bars represent SEM.

